# S-PLM-TM enables fast and accurate sequence-based search for remote homologs

**DOI:** 10.64898/2026.11.16.767046

**Authors:** Duolin Wang, Yichuan Zhang, Shuai Zeng, Yuexu Jiang, Qing Shao, Dong Xu

## Abstract

Protein structures and functions can remain conserved after extensive sequence divergence, but detecting these distant relationships across large databases remains challenging. Here we present S-PLM-TM, a sequence-only remote homology search tool based on a structure-aware protein language model. Cross-modal flow matching and contrastive learning incorporate structural information into sequence representations. A lightweight calibration head enables structural-similarity search with the frozen encoder, while a residue-level alignment head generates structural alignments directly from sequence. S-PLM-TM outperforms the tested sequence-based methods in remote homology detection, rivals structure-based methods in structural classification and retrieves the most functionally consistent neighbors. It also retrieves distantly related enzymes with shared catalytic functions at a level comparable to the best-performing structure-based methods. In large-scale database search, S-PLM-TM achieves the highest average precision among both sequence-based and alignment-free methods, with speed approaching the fastest sequence-alignment tools. Together, these results establish S-PLM-TM as a practical tool that combines sensitive and biologically informative remote homology search with the speed required for database-scale retrieval.

## Introduction

Sequence homology search is a fundamental operation in molecular biology. Tools such as BLAST and PSI-BLAST^1^, HMMER^2^, HHblits^3^, MMseqs2^4^ and DIAMOND^5^ enable functional annotation of newly identified proteins^6^, generate multiple sequence alignments for protein structure prediction^7,8^, provide evolutionary constraints for protein design^9^ and trace the evolutionary history of protein families. However, their sensitivity and specificity decline with sequence similarity. Once pairwise identity falls below 25– 30%, the so-called twilight zone, sequence alignment can no longer tell true homologs from unrelated proteins^10^. Consequently, many proteins identified in newly sequenced genomes and metagenomes cannot be linked to characterized relatives and remain functionally unannotated^11^. Protein structure, however, is conserved three to ten times more than sequence^12,13^. It allows remote homologs to retain common folds, common ancestors, and related functions where their sequences have diverged beyond recognition. With more than 200 million predicted structures now available^14^, structure-based comparisons have uncovered distant relationships missed by conventional sequence search^15^. The challenge is to recover this structural signal from sequence alone with sufficient sensitivity, fast enough to search databases of hundreds of millions of proteins.

Protein language models (PLMs), trained on hundreds of millions of unlabeled sequences, offer another approach to this problem^16–18^. Their representations encode evolutionary and structural regularities that substitution matrices cannot capture, and they have been exploited in two complementary ways. One line uses per-residue embeddings as a learned substitution score inside dynamic-programming alignment, as in pLM-BLAST^19^, embedding-based alignment^20^ and related approaches^21^. The other dispenses with residue-by-residue alignment and compares whole-protein representations, treating proximity in embedding space as a soft, alignment-free measure of relatedness^22^. Because each protein becomes a single vector, search reduces to a nearest-neighbor lookup that can be indexed and scaled to entire sequence databases.

Representation-based search methods differ mainly in how structural knowledge reaches the representation. A first family builds on a pretrained PLM and learns, from structural alignments, a mapping onto structural similarity. TM-Vec^23^ places a twin network on top of ProtT5 embeddings^18^ and trains it to regress the TM-scores computed by TM-align^24^ for millions of protein pairs, so that cosine similarity approximates TM-score; its companion model, DeepBLAST, learns structural alignments from the same backbone. PLMSearch^25^ follows the same logic on ESM-1b embeddings^16^, training a similarity predictor on TM-scores of SCOPe40 and CATH S40 pairs, adding a Pfam clan^26^ prefilter for speed and re-ranking candidates with embedding-based alignment (PLMAlign). A second family treats search as an end-to-end retrieval system. DHR^27^ initializes a dual encoder from ESM, trains it contrastively on homologous pairs drawn from multiple sequence alignments and retrieves homologs from tens of millions of sequences in seconds, whereas ERAST^28^ couples ESM-2 embeddings to a billion-scale vector database with HMMER-based pre-retrieval and post-retrieval refinement. In all of these methods, however, the encoder itself never observes a three-dimensional structure. Structure enters, at most, as a scalar label attached to protein pairs, and the underlying embeddings are shaped by masked-language-model pretraining that is dominated by sequence-level statistics, precisely the signal that vanishes in the remote-homology regime.

When structures are available, structure comparison remains the most sensitive way to detect remote homologs. Alignment-based tools such as TM-align^24^, Dali^29^ and GTalign^30^ superpose three-dimensional coordinates directly, GTalign using spatial indexing to accelerate the search, whereas Foldseek^31^ encodes tertiary interactions into a structural alphabet (3Di) so that fast sequence-search machinery can be applied to structures. Embedding-based tools such as GraSR^32^ and FoldExplorer^33^ enable fast vector-based searches by learning protein representations from residue contact graphs, with FoldExplorer additionally incorporating ESM-2 sequence features. All of them, however, require a reliable structure for the query and for every database entry. For most sequences, including metagenomic proteins, designed variants, newly sequenced genomes, such structures are not available, and predicting them adds a computational step that dwarfs the search itself. What is missing is a method that carries structural knowledge inside a sequence-only representation.

Our previous structure-aware PLM (S-PLM) represented initial steps towards this goal, using sequence–structure contrastive learning to encode three-dimensional structural information into sequence-only representations^34,35^. Here we introduce S-PLM-TM, a new architecture for structure-aware sequence representation learning, and apply it to database-scale remote homology search. During training, S-PLM-TM couples sequence and structure encoders through complementary learning objectives: residue-level cross-modal flow matching^36^ maps sequence representations onto their structural counterparts, whereas protein-level contrastive learning with structure-softened negatives organizes protein representations according to structural similarity. All objectives act in continuous representation space, so the sequence model is never required to predict structure explicitly, whether as structural tokens or as coordinates, and the pretrained PLM remains frozen apart from lightweight adapters. At inference, only the sequence encoder is used. A lightweight calibration head, trained on measured TM-scores with the encoder frozen, then converts embedding similarity into a structural-similarity score for database-scale retrieval. As an additional capability, a residue-level head trained with TM-align supervision^37^ generates sequence-based alignments for retrieved candidates. Because the encoder remains frozen, task-specific heads can be trained on the embedding produced by the calibration head without retraining it; we use one such head for Enzyme Commission (EC) retrieval.

Across complementary evaluations of evolutionary relatedness^38^, structural classification^39^ and molecular function, S-PLM-TM improves sequence-only remote homology search. It leads tested sequence-based methods in the sequence-identity twilight zone and approaches or exceeds methods requiring three-dimensional structures in structural classification and functional consistency. The enzyme-retrieval results extend this advantage to shared catalytic function across divergent sequences, achieving performance comparable to leading structure-based methods, including Dali and Foldseek. At the database scale, S-PLM-TM achieves the highest average precision among the evaluated sequence-based and alignment-free methods while approaching the speed of the fastest sequence-alignment tools. A residue-level head additionally recovers structural correspondences directly from sequence, including those between structurally analogous proteins^40^. These results indicate that encoding structural knowledge in sequence-only representations brings structure-level sensitivity to remote homology search with the speed required for large-scale retrieval.

## Results

### Overview of S-PLM-TM

S-PLM-TM is a structure-aware sequence-search framework that uses amino-acid sequences alone to retrieve remote homologs, estimate their structural similarity and align selected hits at the residue level (Fig. 1). During pretraining, paired amino-acid sequences and protein backbones are processed by sequence (ProtT5-XL^18^) and structure encoders (GVP^41^), respectively. Cross-modal flow matching aligns their representations at the residue and protein levels, complemented by protein-level contrastive learning that attracts matched pairs and repels different proteins using structure-softened negatives (Fig. 1a). Together, these objectives incorporate structural information into the sequence representations.

**Figure 1.**
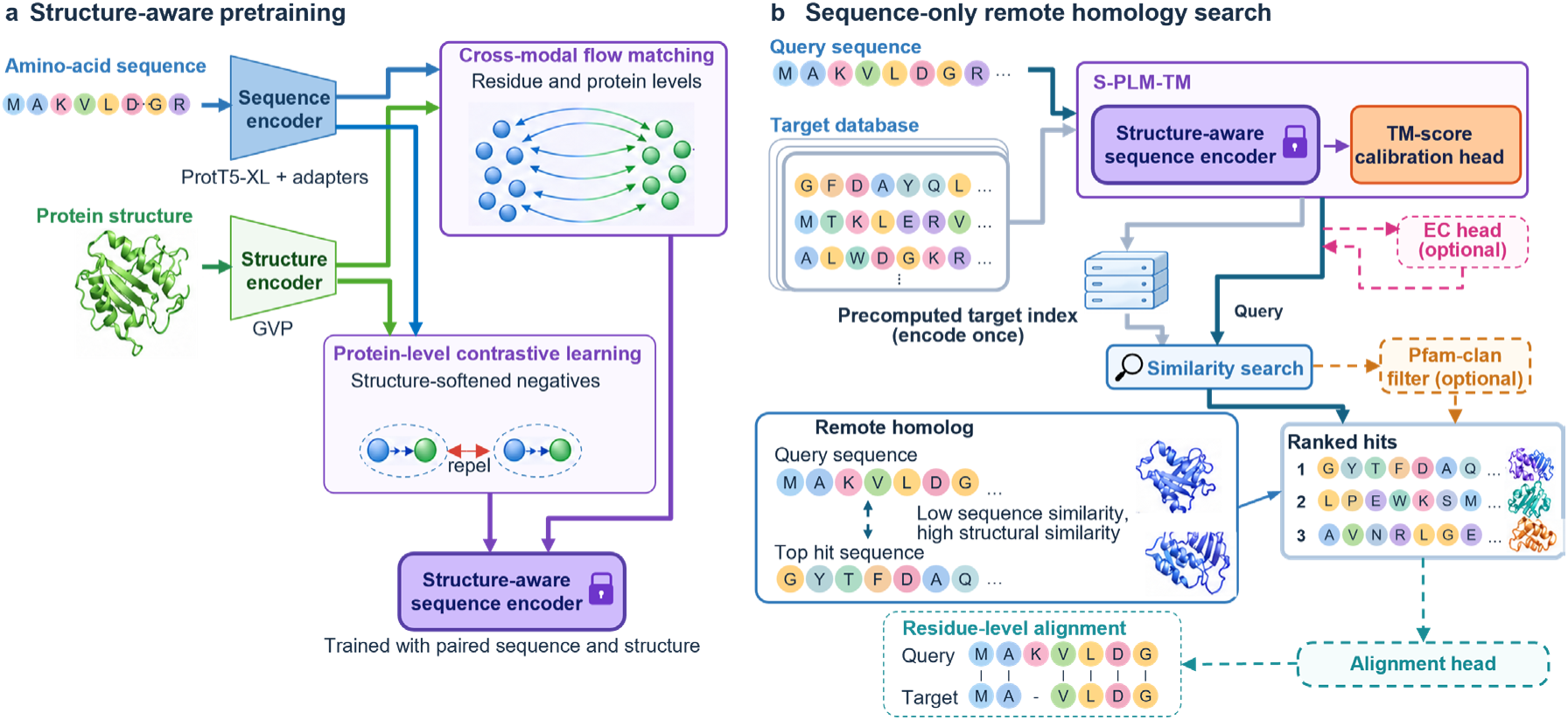
Overview of S-PLM-TM for sequence-only remote homology search. ***a,*** Structure-aware pretraining integrates a ProtT5-XL sequence encoder with trainable adapters and a GVP structure encoder. Cross-modal flow matching aligns sequence and structure representations at the residue and protein levels. Protein-level contrastive learning attracts matched pairs and repels different proteins using structure-softened negatives. ***b***, For sequence-only remote homology search, S-PLM-TM combines the frozen structure-aware sequence encoder with a TM-score calibration head. Query and target sequences are encoded independently, with target representations precomputed once and indexed. An optional EC head produces EC-specific embeddings, while an optional Pfam-clan filter restricts ranking to clan-matched targets when available and otherwise retains full-database ranking. Similarity-based ranking can place structurally similar targets among the top hits even when sequence similarity is low. An alignment head generates residue-level alignments for selected query–target pairs.

For sequence-only remote homology search, the frozen structure-aware sequence encoder is coupled to a lightweight TM-score calibration head trained with TM-align supervision (Fig. 1b). The head maps protein-level representations into calibrated embeddings whose cosine similarity estimates structural similarity. Throughout this study, S-PLM-TM refers to this encoder-plus-head system, analogous to TM-Vec and PLMSearch, which likewise combine a pretrained protein language model with a learned similarity module (Methods). Query and target sequences are encoded independently, with target embeddings computed once and indexed. Queries are compared against this precomputed index without requiring predicted structures for either side (Fig. 1b). Similarity-based ranking can place structurally similar targets among the top hits even when sequence similarity is low. An optional Pfam-clan filter adapted from PLMSearch²⁵ restricts ranking to targets sharing at least one clan with the query when such matches are available; otherwise, full-database ranking is retained. This filter provides a coarse evolutionary constraint, while S-PLM-TM embedding similarity determines the ordering of retained targets (Methods).

Beyond ranking targets by structural similarity, two optional heads support enzyme-specific retrieval and residue-level alignment (Fig. 1b). Enzyme retrieval requires greater functional specificity than what global structural similarity alone can provide, because catalytic function is often determined by a few active-site residues and can differ among proteins sharing a fold^42–44^. To address this, a dedicated EC head trained on EC annotations transforms TM-calibrated protein representations into EC-specific embeddings. Separately, an alignment head converts residue-level sequence representations from selected query–target pairs into a residue–residue similarity matrix that supports both global (Needleman–Wunsch^45^) and local (Smith–Waterman^46^) alignment (Methods).

We evaluated S-PLM-TM on four benchmarks spanning remote homology detection, fold classification and functional retrieval, followed by a large-scale assessment of search accuracy and speed. Comparisons included both sequence-based methods and structure-based aligners, with their different input requirements clearly indicated.

### S-PLM-TM leads every sequence-based method in the twilight zone

Remote homology search is particularly challenging in the sequence-identity twilight zone, defined here as below 30% identity, where sequence similarity alone provides limited evidence of common evolutionary ancestry^10^. To evaluate retrieval in this regime, we used SCOPe protein domains as queries and database targets. SCOPe classifies domains by structural and evolutionary relationships: families contain close relatives, whereas superfamilies unite families for which structural and functional evidence supports a common ancestor^38^. This hierarchy provides reference homology assignments beyond readily detectable sequence similarity.

We first assessed family- and superfamily-level retrieval in 10-percentage-point identity bins using AUC-1FP, the fraction of relevant domains retrieved before the first false positive (Fig. 2a, b; Methods). Across the three bins below 30% identity, S-PLM-TM achieved the highest mean AUC-1FP at both family and superfamily levels, outperforming all eleven baselines in each comparison (Fig. 2a, b). At the family level, it reached AUC-1FP 0.482, 0.788 and 0.942 in the [0,10), [10,20) and [20,30)% bins, against 0.419, 0.717 and 0.886 for PLMSearch and 0.252, 0.698 and 0.909 for TM-Vec, the strongest sequence-based alternatives. The advantage extended to superfamily-level retrieval: S-PLM-TM reached 0.560, 0.763 and 0.926 against 0.504, 0.717 and 0.889 for PLMSearch. We assessed the statistical significance of these differences using per-query paired tests (Methods). Against the strongest alternative in each bin, S-PLM-TM showed significant improvements in the [10,20) and [20,30)% bins at both family level (adjusted *P* = 2.3 × 10⁻¹⁸ and 2.5 × 10⁻⁶, respectively) and superfamily level (adjusted *P* = 2.5 × 10⁻⁶ and 6.7 × 10⁻⁶, respectively). In the sparsely represented [0,10)% bin, comprising only 59 family-level and 94 superfamily-level queries, S-PLM-TM also achieved the highest mean AUC-1FP: 0.482 and 0.560, respectively, compared with 0.419 and 0.504 for PLMSearch (adjusted *P* = 0.050 and 0.12). The differences narrowed at high sequence identity: in the [90,100]% bin, the full spread across all methods was 0.002 at family level (Supplementary Fig. 1).

**Figure 2.**
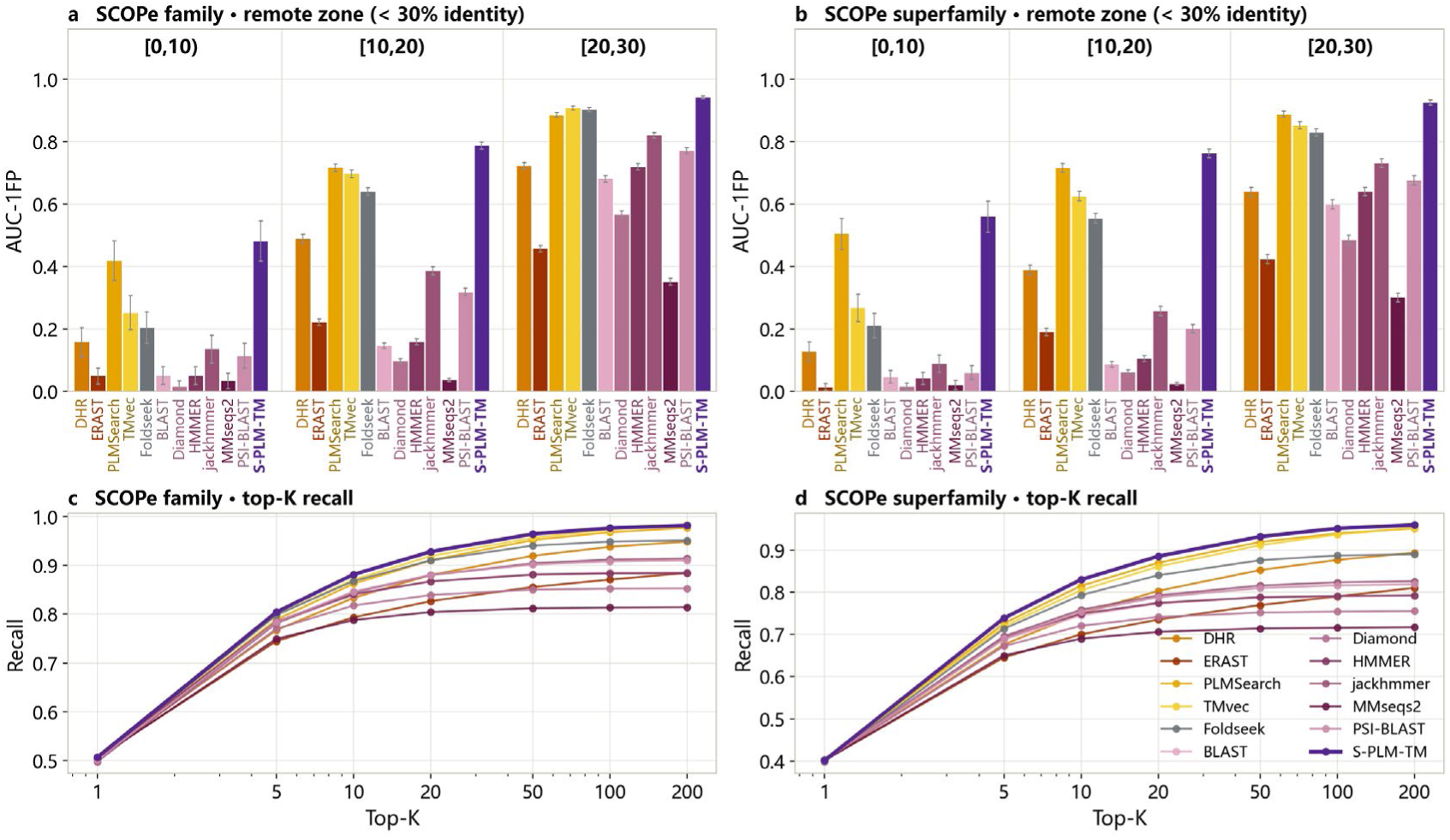
Remote homology detection on SCOPe. **a, b**, Mean AUC-1FP ± s.e.m. in the three sequence-identity bins below 30% for family-level (a) and superfamily-level (b) retrieval. Bars are grouped by identity bin. **c, d**, Top-K recall across the full sequence-identity range for family-level (c) and superfamily-level (d) retrieval.

We next assessed top-K recall across the full sequence-identity range, measuring the fraction of same-family or same-superfamily targets recovered among each query’s K highest-ranked hits, averaged across queries (Fig. 2c, d). At the family level, recall reached 0.881 at K = 10 and 0.976 at K = 100, against 0.871 and 0.973 for TM-Vec. At the superfamily level, the corresponding values were 0.830 and 0.951, against 0.815 and 0.939 for PLMSearch. Thus, S-PLM-TM improved remote homology detection in the twilight zone and recovered more homologs among the top-ranked hits.

### Fold classification approaches structure-based methods

Protein fold space is organized hierarchically, from broad structural classes to homologous superfamilies that capture more remote evolutionary relationships. We therefore evaluated S-PLM-TM on the CATH-S20^39^ fold-classification task included in a recent benchmark^47^ that assesses retrieval across four structural levels, using 15,043 domains (Fig. 3; Methods). In the two-dimensional projection of the S-PLM-TM sequence embeddings, domains from the same class occupy distinct regions, whereas homologous superfamilies form compact and well-separated clusters (Fig. 3a).

**Figure 3.**
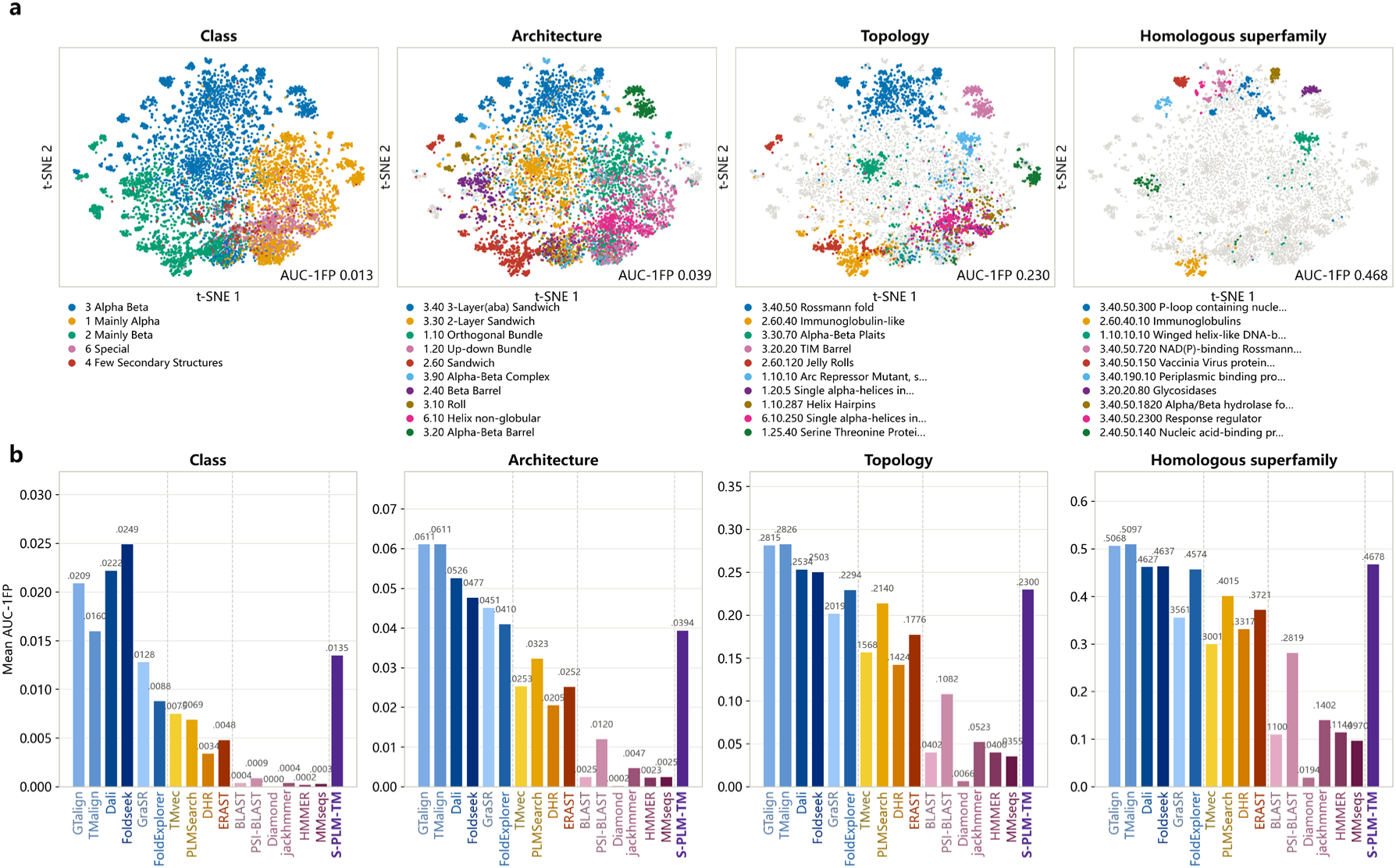
Fold classification on CATH-S20. **a**, Two-dimensional t-SNE projection of the S-PLM-TM sequence embeddings of all 15,043 CATH-S20 domains. The four panels show the same projection colored by the largest categories at the class, architecture, topology and homologous-superfamily levels (all five classes; the ten largest categories at the other three levels); domains outside those categories are grey. The AUC-1FP value for the corresponding level is given in each panel. **b**, Mean AUC-1FP of S-PLM-TM and sixteen published methods at the same levels, computed over the complete ranking of all targets. Methods are ordered by the input they consume, with S-PLM-TM shown last; dashed lines separate structure-based, representation-based sequence-search and alignment-based sequence-search methods. All baselines were reproduced in house under the same CATH domain annotations and evaluation code.

We evaluated S-PLM-TM alongside sixteen published baselines, grouped by the type of input they consume (Fig. 3b). The relative ordering obtained in-house was consistent with that reported in the original benchmark. Measured as AUC-1FP, S-PLM-TM obtained 0.013, 0.039, 0.230 and 0.468 at the class, architecture, topology and homologous-superfamily levels, respectively, and was the highest-scoring sequence-based method at every level (adjusted *P* ≤ 7.9 × 10⁻²¹ against the strongest sequence-based method at each level). The margin was widest at the homologous-superfamily level, the level that corresponds to remote homology: 0.468 against 0.402 for PLMSearch and 0.300 for TM-Vec, the strongest sequence-based alternatives, placing S-PLM-TM third overall behind only TM-align (0.510) and GTalign (0.507). Structural aligners stayed ahead at the three coarser levels, where recovering an entire class or architecture favors methods that compare structures directly. Mean F1-max, which rewards deep recall rather than the top of the ranking, placed S-PLM-TM first among sequence-based methods at the architecture and topology levels, with PSI-BLAST higher at the class level and PLMSearch at the homologous-superfamily level (Supplementary Fig. 2).

We compared S-PLM-TM with the two earlier S-PLM versions on the same benchmark (Supplementary Fig. 3). Performance improved across versions at both the topology and homologous-superfamily levels: AUC-1FP increased from 0.136 to 0.155 and 0.230 at the topology level, and from 0.308 to 0.347 and 0.468 at the homologous-superfamily level. Mean F1-max showed the same ordering. The TM-score calibration head further improved fine-grained retrieval. Adding the head increased AUC-1FP from 0.186 to 0.230 at the topology level and from 0.386 to 0.468 at the homologous-superfamily level, while reducing the class-level score from 0.032 to 0.013. Consistent with this trade-off, the embedding projection showed less contiguous class-level regions but tighter and better-separated homologous-superfamily clusters (Supplementary Fig. 3c, d).

### Retrieved neighbors are the most functionally consistent

Retrieving proteins with related biochemical functions provides a basis for annotating uncharacterized sequences. Using the functional-consistency task of the same benchmark^47^, we compared Gene Ontology^48^ Molecular Function (GO-MF) annotations for each query and its top-10 retrieved neighbors using semantic similarity (Fig. 4a, b; Methods). Across all annotated query–target pairs, S-PLM-TM obtained a mean GO-MF similarity of 0.572 ± 0.003 (mean ± standard error of the mean, i.e., s.e.m.), the highest of all seventeen methods, ahead of DHR (0.556 ± 0.003) and the structural aligners Dali (0.553 ± 0.003) and TM-align (0.541 ± 0.003); paired by query, the margin over DHR was significant (adjusted *P* = 5.0 × 10^−4^). S-PLM-TM also had the highest median score (0.590), close to its mean, whereas HMMER and MMseqs had substantially lower medians than means (0.262 versus 0.381 and 0.292 versus 0.404, respectively), consistent with right-skewed distributions (Fig. 4a, b).

**Figure 4.**
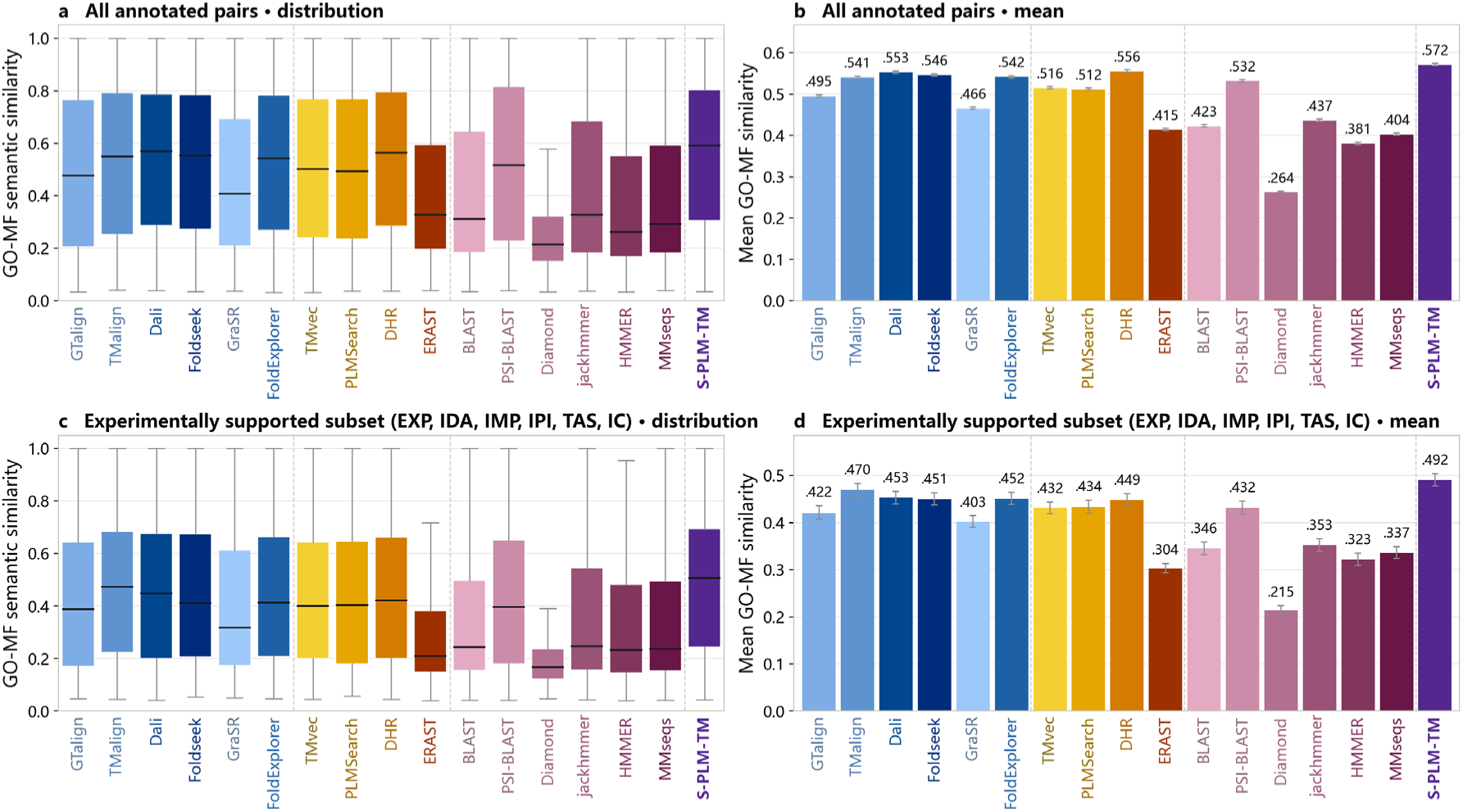
Functional consistency of retrieved neighbors. Gene Ontology Molecular Function (GO-MF) semantic similarity between a query and each of its ten highest-ranked targets, scored as a best-match average of GO-term similarities. **a, b**, All annotated query–target pairs. **c, d**, The experimentally supported subset. **a, c**, Boxes span the interquartile range and the horizontal line marks the median; whiskers extend to the most extreme values within 1.5 times the interquartile range of the box edges. Outliers are not shown. **b, d**, Mean ± s.e.m. across query–target pairs. Dashed lines separate structure-based, representation-based sequence-search and alignment-based sequence-search methods.

To reduce dependence on computationally inferred function labels, we repeated the evaluation on the experimentally supported subset defined by the evidence codes of the published benchmark. S-PLM-TM achieved the highest observed mean score, reaching 0.492 ± 0.013, compared with 0.470 ± 0.013 for TM-align, 0.453 ± 0.014 for Dali and 0.449 for DHR, the strongest sequence-based alternative (Fig. 4c, d; adjusted *P* = 0.37, 0.12 and 0.037, respectively). This smaller subset included 150 queries with experimentally supported annotations on both sides.

### Sequence-only retrieval of microbial–host isozymes

Identifying bacterial enzymes that catalyze the same reactions as human enzymes helps connect gut microbial metabolism to host physiology. This microbial–host isozyme retrieval task requires recognizing shared catalytic function across highly divergent sequences. We used the dataset curated in the DEER study^49^, which selected EC families with an average sequence identity below 10% between bacterial and eukaryotic counterparts. Human enzyme templates were searched against 4,213 bacterial enzymes, including 449 known isozyme targets, with a correct hit defined by a shared EC number (Fig. 5; Methods).

**Figure 5.**
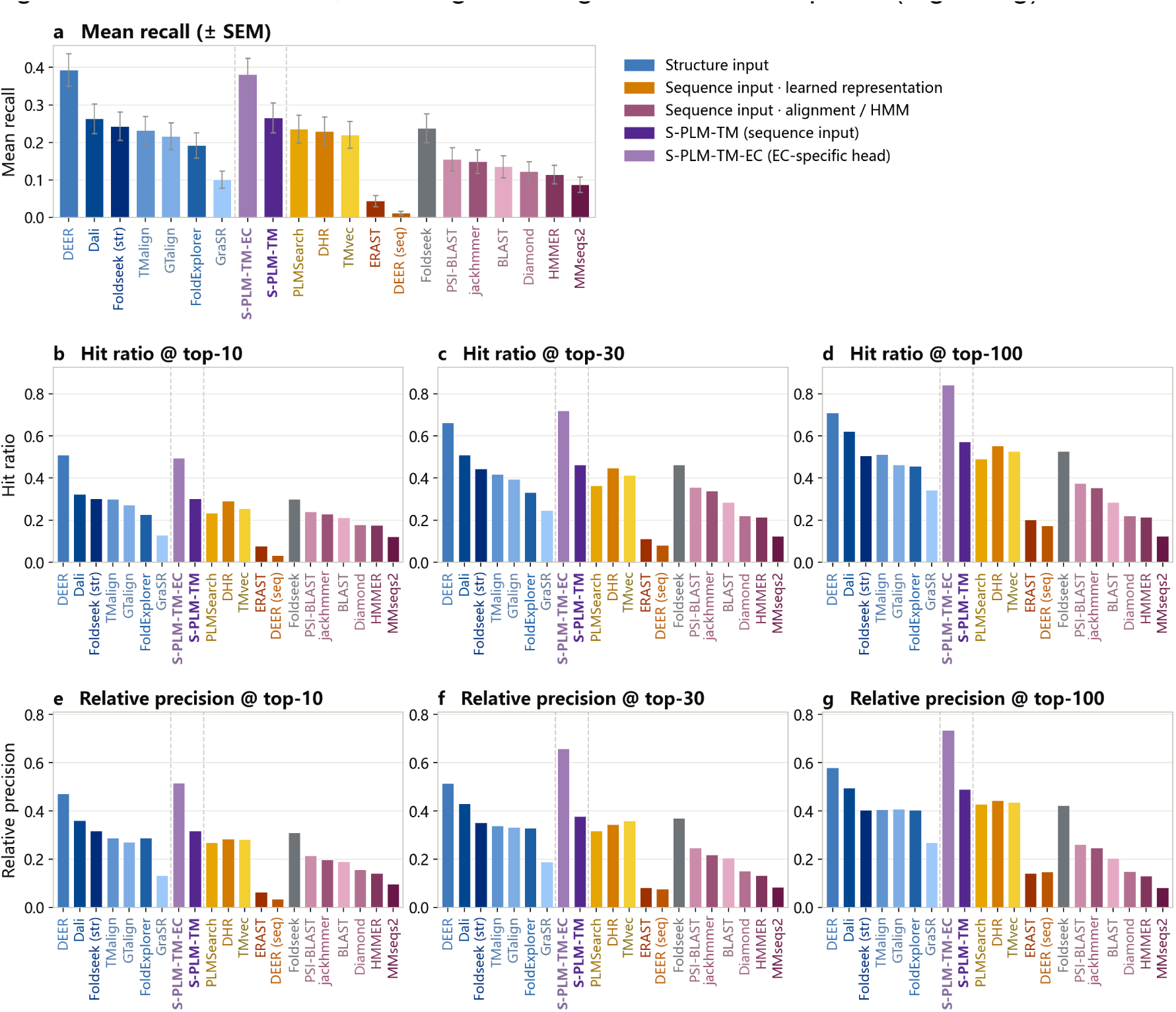
Microbial–host isozyme retrieval. Human enzymes serve as templates to retrieve their bacterial counterparts from a curated set of 4,213 bacterial enzymes containing 449 known isozyme targets; a retrieved enzyme counts as correct when it carries the template’s EC number. **a**, Mean recall: for a template with *n* true counterparts, the fraction of those *n* recovered within its top *n* hits, averaged over templates; error bars show the s.e.m. over templates. **b**–**d**, Hit ratio: the fraction of the 449 true isozymes recovered by at least one template within its top 10, 30 or 100 hits. **e**–**g**, Relative precision: correct hits among the top *k* divided by min(*k*, *n*), averaged over templates, so that 1 means every attainable counterpart is ranked within the cut-off. Bars are colored by the input a method consumes and ordered by mean recall within each group; dashed lines mark the group boundaries. S-PLM-TM-EC, S-PLM-TM with the EC-specific retrieval head; Foldseek (str), Foldseek in structure mode; DEER (seq), DEER’s sequence-only ablation.

On the 90 EC-family templates released with DEER, one representative per family, S-PLM-TM reached a mean recall of 0.266 ± 0.040 (mean ± s.e.m.), the highest of the eighteen methods evaluated without EC-level supervision: ahead of every sequence-based method, including Foldseek in sequence mode (0.238 ± 0.039), PLMSearch (0.236 ± 0.038), DHR (0.230 ± 0.038) and TM-Vec (0.220 ± 0.035), and level with Dali (0.263 ± 0.040), the strongest structural aligner here (Fig. 5a). At the top-100 cut-off its relative precision of 0.491 was second only to Dali (0.496; Fig. 5g).

To assess EC-specific retrieval, we trained the optional EC head on EC annotations while keeping the encoder and TM-score calibration head frozen (Methods). Under the same protocol, this sequence-only model reached a mean recall of 0.382±0.042 against 0.393 ± 0.043 for the structure-input DEER model (Fig. 5a); the difference of 0.011 was not significant (adjusted *P*=0.73). The margin over the strongest sequence-based alternative, Foldseek in sequence mode (0.238), was significant (adjusted P=4.3 × 10^−4^), and S-PLM-TM was ahead of every other method tested except DEER (adjusted P≤1.8 × 10^−3^). At top 30 and top 100, it recovered 0.719 and 0.842 of the 449 positive bacterial targets, compared with 0.662 and 0.708 for DEER (Fig. 5c, d). Relative precision was higher at all three cut-offs, reaching 0.736 against 0.580 at top 100 (Fig. 5e–g).

### Search at scale, and residue-level structural alignment

To assess sequence-only remote homology search at database scale, we assembled 957 SCOPe query sequences and searched them against 430,140 Swiss-Prot target sequences. TM-align scores between each query and target provided the structural ground truth; query selection, target construction and evaluation are described in Methods.

S-PLM-TM achieved a mean average precision (MAP) of 0.638 over all positives and 0.620 among positives below 30% sequence identity, the highest of any sequence-based methods. It outperformed every method that uses structure as input except TM-align and GTalign. Precision reached 0.844 at rank 1 (P@1) and 0.805 among the top ten hits (P@10) (Table 1). TM-align was excluded from statistical comparisons because it defines the ground truth. Across the 957 queries, S-PLM-TM significantly outperformed all remaining methods except GTalign in MAP, MAP (<30%) and P@10 (adjusted P ≤ 6.4 × 10⁻⁸, ≤ 1.3 × 10⁻⁸ and ≤ 0.015, respectively). Compared with the closest competitor, TM-Vec, the gains were 0.035 in MAP, 0.039 in MAP (<30%) and 0.028 in P@10 (adjusted *P* = 6.3 × 10⁻⁸, 9.6 × 10⁻⁹ and 1.3 × 10⁻⁶, respectively). P@1 was 0.844 for S-PLM-TM and 0.829 for TM-Vec (adjusted *P* = 0.098). Within sequence-identity bins, S-PLM-TM ranked first among sequence-based methods at both 10–20% (MAP 0.569) and 20–30% (MAP 0.665) identity (Supplementary Table 1). S-PLM-TM achieved this without any Pfam prefilter, whereas PLMSearch applies its own Pfam-clan prefilter and jackhmmer and PSI-BLAST build iterative sequence profiles, so S-PLM-TM ranked first among sequence-based methods while using the least prior information. Applying the same Pfam-clan prefilter to S-PLM-TM left this ordering unchanged, with a MAP of 0.601 against 0.580 for PLMSearch (two-sided Wilcoxon signed-rank *P* = 1.7 × 10⁻⁶), even though the prefilter lowered its own MAP (Supplementary Table 5).

**Table 1.**
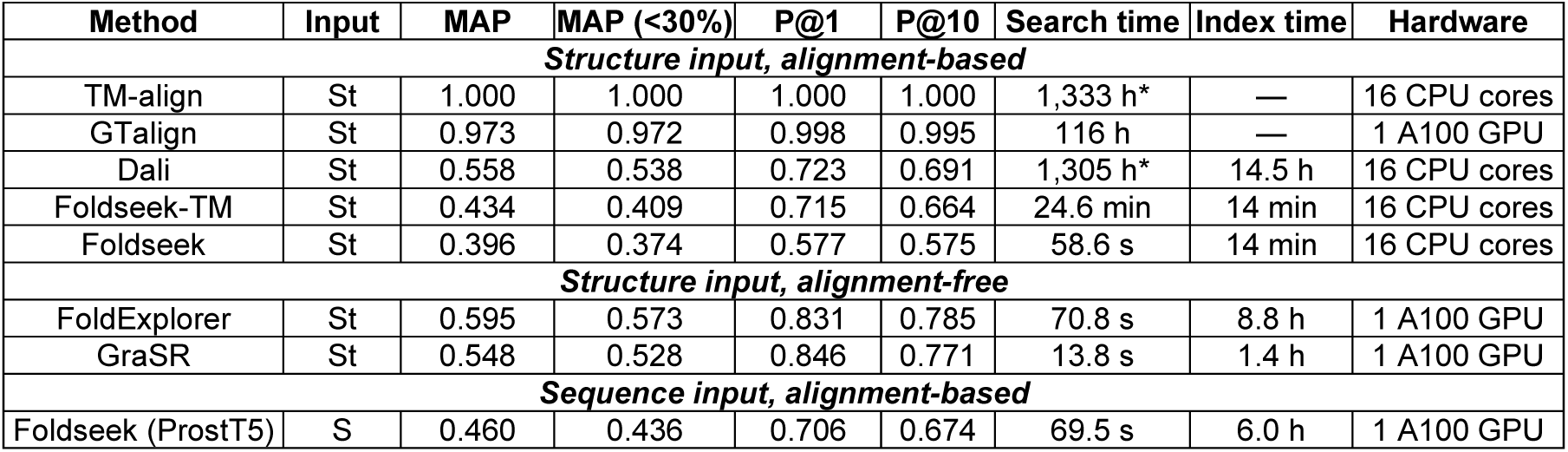
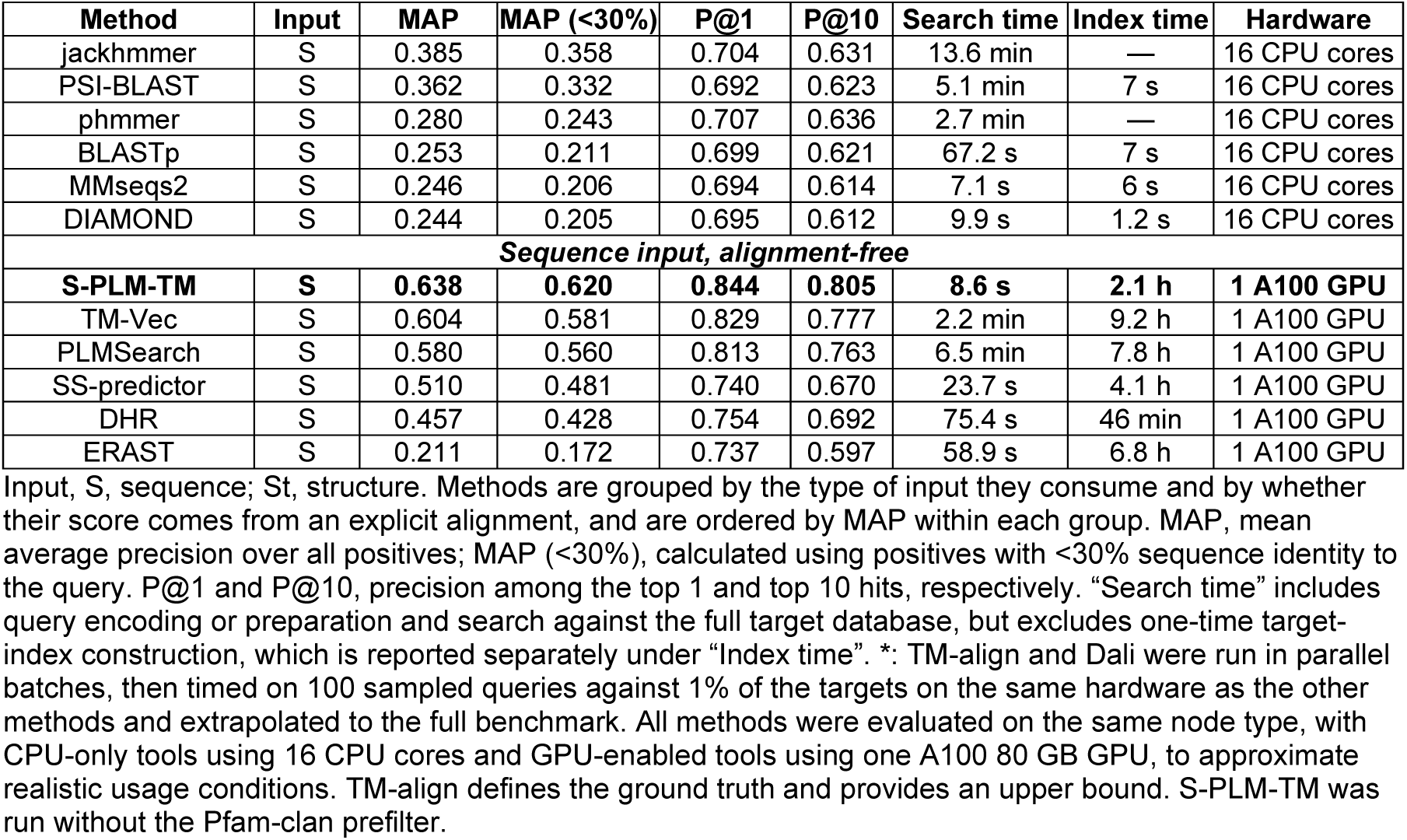
Search performance and computational cost at scale.

| Method | Input | MAP | MAP (<30%) | P@1 | P@10 | Search time | Index time | Hardware |
|---|---|---|---|---|---|---|---|---|
| *Structure input, alignment-based* |  |  |  |  |  |  |  |  |
| TM-align | St | 1.000 | 1.000 | 1.000 | 1.000 | 1,333 h* | — | 16 CPU cores |
| GTalign | St | 0.973 | 0.972 | 0.998 | 0.995 | 116 h | — | 1 A100 GPU |
| Dali | St | 0.558 | 0.538 | 0.723 | 0.691 | 1,305 h* | 14.5 h | 16 CPU cores |
| Foldseek-TM | St | 0.434 | 0.409 | 0.715 | 0.664 | 24.6 min | 14 min | 16 CPU cores |
| Foldseek | St | 0.396 | 0.374 | 0.577 | 0.575 | 58.6 s | 14 min | 16 CPU cores |
| *Structure input, alignment-free* |  |  |  |  |  |  |  |  |
| FoldExplorer | St | 0.595 | 0.573 | 0.831 | 0.785 | 70.8 s | 8.8 h | 1 A100 GPU |
| GraSR | St | 0.548 | 0.528 | 0.846 | 0.771 | 13.8 s | 1.4 h | 1 A100 GPU |
| *Sequence input, alignment-based* |  |  |  |  |  |  |  |  |
| Foldseek (ProstT5) | S | 0.460 | 0.436 | 0.706 | 0.674 | 69.5 s | 6.0 h | 1 A100 GPU |
| jackhmmer | S | 0.385 | 0.358 | 0.704 | 0.631 | 13.6 min | — | 16 CPU cores |
| PSI-BLAST | S | 0.362 | 0.332 | 0.692 | 0.623 | 5.1 min | 7 s | 16 CPU cores |
| phmmer | S | 0.280 | 0.243 | 0.707 | 0.636 | 2.7 min | — | 16 CPU cores |
| BLASTp | S | 0.253 | 0.211 | 0.699 | 0.621 | 67.2 s | 7 s | 16 CPU cores |
| MMseqs2 | S | 0.246 | 0.206 | 0.694 | 0.614 | 7.1 s | 6 s | 16 CPU cores |
| DIAMOND | S | 0.244 | 0.205 | 0.695 | 0.612 | 9.9 s | 1.2 s | 16 CPU cores |
| *Sequence input, alignment-free* |  |  |  |  |  |  |  |  |
| **S-PLM-TM** | **S** | **0.638** | **0.620** | **0.844** | **0.805** | **8.6 s** | **2.1 h** | **1 A100 GPU** |
| TM-Vec | S | 0.604 | 0.581 | 0.829 | 0.777 | 2.2 min | 9.2 h | 1 A100 GPU |
| PLMSearch | S | 0.580 | 0.560 | 0.813 | 0.763 | 6.5 min | 7.8 h | 1 A100 GPU |
| SS-predictor | S | 0.510 | 0.481 | 0.740 | 0.670 | 23.7 s | 4.1 h | 1 A100 GPU |
| DHR | S | 0.457 | 0.428 | 0.754 | 0.692 | 75.4 s | 46 min | 1 A100 GPU |
| ERAST | S | 0.211 | 0.172 | 0.737 | 0.597 | 58.9 s | 6.8 h | 1 A100 GPU |

Input, S, sequence; St, structure. Methods are grouped by the type of input they consume and by whether their score comes from an explicit alignment, and are ordered by MAP within each group. MAP, mean average precision over all positives; MAP (<30%), calculated using positives with <30% sequence identity to the query. P@1 and P@10, precision among the top 1 and top 10 hits, respectively. “Search time” includes query encoding or preparation and search against the full target database, but excludes one-time target-index construction, which is reported separately under “Index time”. \*: TM-align and Dali were run in parallel batches, then timed on 100 sampled queries against 1% of the targets on the same hardware as the other methods and extrapolated to the full benchmark. All methods were evaluated on the same node type, with CPU-only tools using 16 CPU cores and GPU-enabled tools using one A100 80 GB GPU, to approximate realistic usage conditions. TM-align defines the ground truth and provides an upper bound. S-PLM-TM was run without the Pfam-clan prefilter.

Search cost varied by orders of magnitude under a common computational environment. CPU-only methods used 16 CPU cores, whereas GPU-enabled methods used one A100 80 GB GPU, allowing each method to run under realistic usage conditions. S-PLM-TM completed all 957 searches in 8.6 seconds, making it the second fastest of the 20 methods, behind only MMseqs2 (7.1 seconds, MAP 0.246). This was 15 times faster than TM-Vec (2.2 minutes), 8 times faster than FoldExplorer (70.8 seconds) and 45 times faster than PLMSearch (6.5 minutes), and roughly five orders of magnitude faster than the alignment-based structural searches, including Dali (1,305 hours) and TM-align (1,333 hours). One-time construction of the S-PLM-TM target index required 2.1 hours, against 9.2 hours for TM-Vec, 8.8 hours for FoldExplorer, which additionally requires a structure for every entry, and 7.8 hours for PLMSearch. Thus, S-PLM-TM combined the strongest sequence-based retrieval performance in this benchmark with leading search speed and modest indexing cost.

Finally, the residue-level alignment head extends the frozen encoder from retrieval to the alignment of retrieved candidates, reading its per-residue representations rather than the calibrated protein-level score. We evaluated the head on two hand-curated structure-alignment sets, MALIDUP^50^ and MALISAM^40^, scoring its alignments under the protocol of CLAlign^37^ so that the values are directly comparable with those it reports. Scored as full-alignment F1 against the curated alignments, S-PLM-TM reached 0.643 on MALIDUP and 0.282 on MALISAM. CLAlign’s models give the strongest results under this protocol: 0.646 and 0.267 for CLAlign-ProtT5, and 0.677 and 0.299 for CLAlign-ProstT5. Our head matches CLAlign-ProtT5 on MALIDUP and exceeds it on MALISAM, the harder set, whose pairs are structurally analogous but not homologous (median sequence identity 8.0%, none above 30%). The gain therefore falls where sequence signal is weakest, the regime that structure-aware pretraining targets. Our alignment head adds only 4.20 million trainable parameters to the frozen encoder. By comparison, each CLAlign model fine-tunes 10.62 million LoRA parameters within a three-billion-parameter backbone (Supplementary Table 2).

### S-PLM-TM leads other protein language models under identical settings

To isolate the contribution of the sequence encoder, we repeated all four benchmarks with the same downstream pipeline. We paired the frozen encoder from each of four models, S-PLM-TM, ProtT5-XL, ProstT5^51^ and ESM2-650M, with a separate TM-score calibration head. All heads used the same training recipe. We applied the Pfam-clan filter where specified by the corresponding benchmark and kept the evaluation protocol unchanged. ProtT5-XL is the backbone retained in S-PLM-TM, so this comparison directly tests the effect of structure-aware pretraining.

S-PLM-TM achieved the best or tied performance across all four benchmarks (Supplementary Table 3). It was highest in every SCOPe bin below 20% sequence identity, highest at every CATH level by both F1-max and AUC-1FP, and highest for GO functional consistency and bacterial EC retrieval at ranks 10 and 30. Differences narrowed in the less challenging regimes: ProstT5 was marginally higher in the 20–30% SCOPe bins, all encoders exceeded 0.97 above 30% identity, and bacterial EC relative precision at rank 100 differed by no more than 0.02. Because the TM-score calibration head, training recipe, filtering and evaluation protocol were matched, the remaining differences primarily reflect the sequence encoder. We assessed the statistical significance of these differences using per-domain and per-query paired tests (Methods). All twenty-four CATH comparisons (four levels × two metrics × three encoders) were significant (adjusted *P* ≤ 3.7 × 10⁻¹²). The SCOPe advantage over ProtT5-XL and ESM2-650M was significant at both levels in the 10–20% and 20–30% identity bins (adjusted *P* ≤ 0.015), and bacterial EC mean recall was significantly higher than ProtT5-XL (adjusted *P* = 0.049) and ProstT5 (adjusted *P* = 0.015). The remaining margins, including those in the 0–10% bins, which contain only 59 family-level and 94 superfamily-level queries, are reported for reference. S-PLM-TM retained its clearest advantage in the remote-homology regime, where sequence identity is lowest. Notably, S-PLM-TM was pretrained on 0.5 million protein structures, compared with approximately 17 million structures used for ProstT5, which learns structure by translating sequences into 3Di structural tokens. Its advantage was therefore observed despite a substantially smaller structural pretraining set.

We next examined which pretraining objective contributed to this advantage. We trained two control encoders with the same ProtT5-XL backbone, data and training schedule as S-PLM-TM (Methods). The first omitted cross-modal flow matching. The second replaced it with mean-squared-error (MSE) regression of sequence representations onto structure representations. Both controls were then calibrated and evaluated exactly as S-PLM-TM (Supplementary Table 4). Removing flow matching had a small but consistent effect. S-PLM-TM was higher at all four CATH levels, and the per-domain advantage was significant at every level (adjusted *P* ≤ 8.1 × 10^−1^^7^), although the differences themselves were small, between 0.0007 and 0.0058. Replacing flow matching with MSE regression was more harmful. Bacterial EC mean recall fell from 0.266 to 0.231 (adjusted *P* = 0.013), and precision at rank 10 also decreased (adjusted *P* = 0.043). Family-level SCOPe AUC-1FP in the 10–20% identity bin decreased slightly, from 0.788 to 0.782 (adjusted *P* = 0.013). GO and the remaining SCOPe bins did not differ significantly. Flow matching therefore incorporated structural information without the loss of functional signal caused by MSE regression.

We then assessed the contributions of the TM-score calibration head and the Pfam-clan prefilter using the same S-PLM-TM encoder (Supplementary Table 4). The effects of the two components varied across benchmarks. Removing the prefilter reduced SCOPe superfamily AUC-1FP in the 0–10% identity bin from 0.560 to 0.175 and the family-level value from 0.482 to 0.225, indicating that clan information aids recognition of highly divergent homologs. This finding supports combining embedding-based retrieval with Pfam-informed candidate selection, as also implemented in PLMSearch²⁵ and ERAST^28^. CATH and GO were unchanged because their pipelines use no prefilter. The effect of the prefilter depends on the target database. On the large-scale search benchmark, where the targets are full-length Swiss-Prot proteins rather than domains, adding the same prefilter lowered MAP instead of raising it, and S-PLM-TM is therefore reported there without it (Supplementary Table 5). Removing the TM-score calibration head reduced CATH homologous-superfamily AUC-1FP from 0.468 to 0.386, GO mean similarity from 0.492 to 0.440 and bacterial EC mean recall from 0.266 to 0.182. These results support the value of TM-score-supervised scoring for structural and functional retrieval, consistent with the use of TM-score supervision in TM-Vec and PLMSearch. Removing both components reduced performance on the SCOPe and bacterial EC retrieval benchmarks.

## Conclusion and discussion

We present S-PLM-TM, a structure-aware protein language model that enables fast and accurate sequence-only remote homology search. We demonstrate its performance across complementary evolutionary, structural and functional benchmarks. For remote homology detection (SCOPe), S-PLM-TM achieved the highest mean AUC-1FP among all tested methods at both family and superfamily levels below 30% sequence identity. For hierarchical structural classification (CATH), it outperformed all tested sequence-based methods in AUC-1FP at every level, approaching the leading structural aligners at the homologous-superfamily level. Its retrieved neighbors also showed the highest mean GO functional similarity among both sequence- and structure-based methods. For microbial–host isozyme retrieval, it matched Dali in mean recall. With the optional EC head, this sequence-only model approached the mean recall of the structure-input DEER model while achieving higher relative precision at every tested cut-off and recovering more known isozymes at ranks 30 and 100. At database scale, S-PLM-TM achieved the highest MAP among sequence-based and alignment-free methods, with search speed approaching the fastest sequence-alignment tools.

Comparisons with other PLMs, including ProtT5-XL, ProstT5 and ESM2-650M, under the same downstream training and evaluation settings highlight the advantages of the S-PLM-TM pretraining model across structural and functional retrieval tasks. Unlike ProstT5, which acquires structural knowledge through translation into discrete structural tokens, S-PLM-TM learns entirely in continuous representation space. It does not require the sequence model to predict structure explicitly, whether as structural tokens or as coordinates. Structure instead shapes the sequence representation as a training signal, so the representation can absorb structural regularities without being reduced to a structural description. This may explain why S-PLM-TM outperformed ProstT5 on both structural and functional retrieval despite far fewer training structures. Its objectives incorporate structural information in complementary ways: protein-level contrastive learning brings paired sequence and structure representations together, while structure-softened negatives reduce the penalty on structurally similar proteins. Flow matching learns a transformation between the two modalities rather than forcing their projected representations to coincide. This less restrictive objective may be appropriate because the structure used for training captures only one conformation and may not contain all of the functional information encoded by the sequence. Among the three objectives evaluated, flow matching achieved the best overall balance between structural and functional retrieval. It achieved higher CATH topology and homologous superfamily AUC-1FP, GO mean similarity and bacterial EC mean recall than either ablation (Supplementary Table 4). The decrease in enzyme retrieval under MSE regression further supports the value of a flexible cross-modal relationship that incorporates structural information while preserving the functional utility of sequence representations.

The Pfam-clan prefilter and TM-score calibration head offer benefits that depend on the retrieval setting. The contribution of the Pfam-clan prefilter depends on what the target database contains. In the SCOPe benchmark both the queries and the targets are single domains, and a query and its true positives usually belong to the same Pfam clan, so the clan constraint retains almost all positives while excluding a large number of unrelated candidates. Its contribution is largest for the most divergent pairs. In a full-length protein database the situation is different, because many remote positives share no Pfam clan with the query. A target excluded by the prefilter is removed from the ranking, so it cannot be retrieved at all. The leading performance in Table 1 therefore does not depend on Pfam annotation. TM-score calibration improves the ranking of fine-grained structural relationships. Similar strategies have also been effective in other retrieval methods, including Pfam-informed search in PLMSearch²⁵ and ERAST^28^ and TM-score-based learning in TM-Vec²³ and PLMSearch²⁵. The optional EC head further adapts these representations to catalytic function, extending retrieval beyond global structural similarity to identify enzymes with shared catalytic activity.

As an alignment-free search tool, S-PLM-TM combines sensitive retrieval with efficient database screening. Representations of proteins in the database can be precomputed and reused, enabling rapid searches from sequence alone without structure prediction or exhaustive pairwise alignment. Its residue-level alignment head further supports the biological interpretation of remote homology search results by identifying alignable regions and their residue correspondences.

Two components of the pipeline are optional, and their utility depends on the search setting. First, the optimal use of the Pfam clan prefilter depends on the target database: the setting that performs best for a domain database is not optimal for a whole proteome database. Users must therefore decide whether to apply the prefilter. One possible alternative is to run each query both with and without the prefilter and combine the two result lists. This strategy could retain close relatives while also recovering candidates that cross clan boundaries. The additional computational cost would be modest, as searching all 957 queries against 430,140 targets takes only 8.6 s. Second, the choice of the EC head depends on the biological objective rather than on the target database. Because this head is trained to identify shared catalytic function, it is most appropriate when the goal is to retrieve enzymes with the same activity. The TM-score calibration head remains the default choice for searches based on structural homology.

Together, these capabilities make S-PLM-TM a practical tool for finding distant relatives and functionally relevant proteins in large sequence databases in seconds. By recovering homologs missed by conventional sequence search, it could expand the sequence diversity available for evolutionary analyses. Further evaluation across a wider range of taxa and database compositions, especially on proteins from underrepresented organisms and metagenomic samples, will help determine how broadly S-PLM-TM generalizes beyond the current benchmarks. Separately, integrating this retrieval capability with multiple sequence alignment could support phylogenetic inference and ancestral sequence reconstruction, helping trace protein evolution and identify ancestral variants for functional characterization.

## Methods

### Paired sequence and structure encoders

S-PLM-TM learns from paired sequences and structures and retrieves remote homologs from sequence alone (Supplementary Fig. 4). The sequence encoder is ProtT5-XL^18^ (24 blocks; width 1,024), with frozen pretrained weights and 1.61 million trainable adapter parameters in its last 12 blocks (Supplementary Fig. 4a). Sequences are truncated to 512 residues during training. For a protein of length *L*, the encoder produces *L* × 1,024 residue features and a 1,024-dimensional protein representation by mean pooling over residues.

Each adapter^52^ is a 1,024 → 64 → 1,024 perceptron with ReLU σ and output layer normalization^53^:

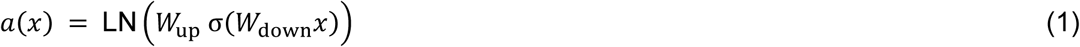

*W*_down_ and *W*_up_ are projection matrices. The residual update scales the adapter output by the root-mean-square (RMS) magnitude of its input and by a learned gain *g*, initialized to 0.44:

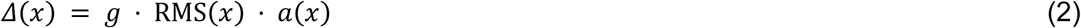

The adapter output is added to the hidden state as x + Δ(x).

The structure encoder is a three-layer GVP^41^ (30 nearest neighbors per residue; node channels of 256 scalars and 32 vectors; edge channels of 32 scalars and one vector), initialized from masked-structure pre-training. It produces *L* × 256 residue features and a 256-dimensional protein representation by mean pooling (Supplementary Fig. 4a).

### Cross-modal representation learning

Cross-modal learning transfers structural information into the sequence representation (Supplementary Fig. 4b). The main objective is flow matching^36^ at the residue and protein levels. Instead of regressing one representation directly onto the other, it learns a velocity field that transports the representation of one modality towards that of the other. At the protein level, contrastive learning with structure-softened negatives organizes proteins by structural similarity. Three supporting objectives stabilize training: masked self-distillation within each modality, variance–covariance regularization and structural auxiliary heads on the structure encoder. All objectives are used only in training. At inference, the sequence encoder alone produces the representation.

Pre-training uses 540,000 Swiss-Prot sequences^54^ paired with AlphaFold backbones^7,14^. Projectors map both modalities to residue (*d* = 64) and protein (*d* = 256) spaces. Online features are trained against stop-gradient exponential-moving-average (EMA) targets^55^ with decay 0.99. Each cross-modal objective is applied in both directions, sequence to structure and structure to sequence (Supplementary Fig. 4b).

For an online source x₀, an EMA target x₁ and *t* sampled uniformly from [0, 1], flow matching^36^ interpolates

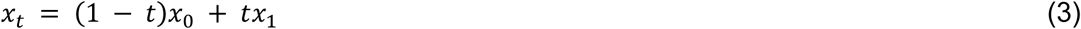

with target velocity v = x₁ − x₀. As in the original formulation^36^, x₀ is scaled by (1 − *σ*_min_) with *σ*_min_ = 10⁻⁵, which is omitted here for clarity. The velocity network is trained with:

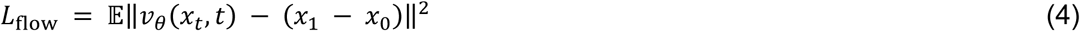

Each direction and representation level has its own velocity network, which takes x_t_ and *t* as input and consists of Linear(*d*, 512), two attention blocks (eight heads) and Linear(512, *d*) (Supplementary Fig. 4b).

### Protein-level contrastive learning

For a batch of B paired proteins, let *h*_1_, …, *h*_2B_ denote the L2-normalized protein-level predictor outputs of the sequence branch and the EMA structure projections, and let *a*⁺ index the cross-modal partner of entry *a*. The InfoNCE loss with structure-softened negatives is

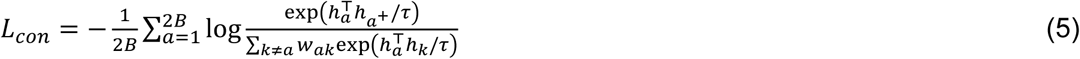

where τ = 0.1, the weight *w*_ak_ = 1 for the positive pair and max(1 − *c*_ak_, 0.01) otherwise, and *c*_ak_ is the cosine similarity between the (detached) structure projections of the proteins at positions *a* and *k*. Structurally similar negatives, which are likely to be remote homologs, therefore contribute little to the denominator, so that the model is not trained to push them apart.

### Within-modality masked self-distillation

Each encoder also predicts its own EMA target from a corrupted input, following BYOL^55^. In the sequence branch, 15% of residues are selected, of which 80% are replaced by a mask token, 10% by a random amino acid and 10% are left unchanged. In the structure branch, 15% of backbone coordinate entries are set to zero, Gaussian noise (s.d. 0.01) is added, or both are applied. Let *f*_θ_ and *f*_ξ_ denote the online and EMA encoder–projector mappings. The online encoder branch (*f*_θ_) receives the corrupted input *x̃*, the target branch (*f*_ξ_) receives the intact input *x*, and a predictor *q* maps the online projection to the target at both the residue and protein levels. At each representation level, we minimize the mean of

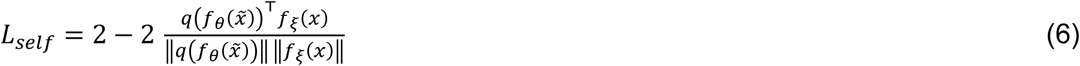

The loss is averaged over the corresponding residue or protein representations.

### Geometry-preserving variance–covariance regularization

At the protein level of each cross-modal direction, the online projection *z*^s^ and the EMA target projection *z*^t^ are L2-normalized and mean-centred over the batch, and a VICReg-style regularizer^56^ is applied without its invariance term:

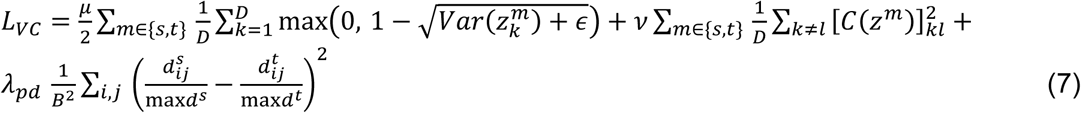

where *C*(·) is the batch covariance matrix, *d*^m^_ij_ is the Euclidean distance between proteins *i* and *j* in modality *m*, D = 256, μ = 25, ν = 1, λ_pd_ = 0.3 and ε is a small constant for numerical stability. The variance and covariance terms discourage collapse and redundant dimensions. The distance term aligns within-batch geometry across modalities. Gradients pass only through the online branch.

### Structural auxiliary heads

Two lightweight heads read from the structure encoder and inject explicit geometric supervision. For the distogram head, up to 256 residue pairs with sequence separation of at least 12 are sampled per protein, and their Cα–Cα distances *d*_ij_ are discretized into 64 bins between 2.3 and 21.7 Å, as in AlphaFold^7^. A perceptron (128 → 256 → 64; ReLU; dropout 0.1) maps the concatenated residue projections [*e*_i_; *e*_j_] to bin probabilities *p*_ϕ_, trained with cross-entropy over the set P of sampled pairs (weight 0.2):

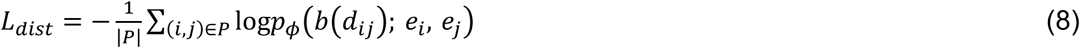

A shape head (256 → 128 → 2; ReLU) predicts, from the protein-level projection, the radius of gyration and the normalized asphericity of the Cα trace,

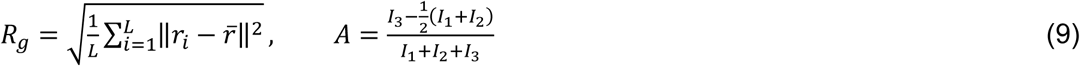

where *r*i are Cα coordinates and *I*1 ≤ *I*2 ≤ *I*3 are the eigenvalues of their inertia tensor. The shape head is trained with a Huber loss (δ = 2; weight 0.1). These heads are attached only to the structure branch; their geometric supervision reaches the sequence representation through the cross-modal objectives.

The total pretraining loss sums the bidirectional flow losses at both representation levels, protein-level contrastive and variance–covariance losses, and within-modality self-distillation losses at both levels. These components have unit outer weights; the internal regularization coefficients and structural auxiliary weights are specified above.

Pre-training was carried out on a cluster over approximately one week, including interruptions, using one A100 GPU at a time, batches of 20 proteins and at most 2,000 residues per batch for residue-level losses. SGD used peak learning rate 0.01, weight decay 5 × 10⁻⁴, gradient clipping at 1, and a 2,000-step cosine cycle with minimum learning-rate ratio 0.01.

### TM-score calibration head

The TM-score calibration head acts on the pooled 1,024-dimensional features of the frozen sequence encoder, before the training projectors (Supplementary Fig. 4c):

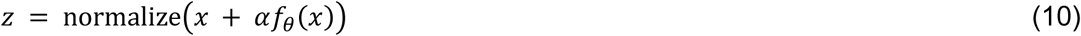

Here *f*θ is a two-layer perceptron, Linear(1,024, 1,024), GELU^57^ and Linear(1,024, 1,024), α is a learned scalar initialized to 1, and normalize denotes L2 normalization (2.10 million parameters). The final layer is zero-initialized.

Training uses 266,500 anchors and 1.41 × 10⁸ measured-neighbor relations from TM-Vec^23^. The target *τ*ij for anchor *i* and measured neighbor *j* is the larger of the two length-normalized TM-align scores^24^. Learned scale *a* and offset *b* calibrate cosine similarity:

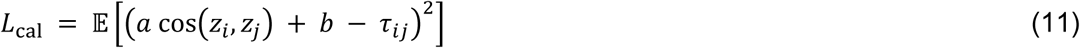

Random pairs from different anchor groups receive a squared hinge penalty:

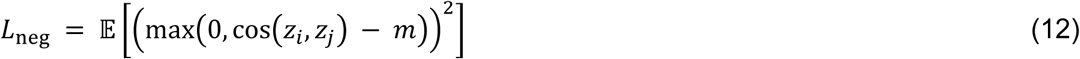

The combined objective is

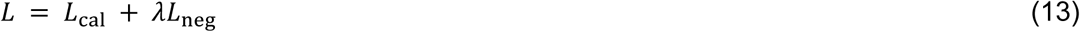

with margin m = 0.3 and λ = 0.1. AdamW^58^ trains the head for 4,000 steps with 2,048 anchors and 16 measured neighbours per anchor, sampled across ten TM-score bins. Head parameters use learning rate 10⁻⁴ and weight decay 10⁻⁴. Validation on 4,096 anchors every 200 steps selects the lowest-MSE checkpoint.

### Optional EC retrieval head

The optional EC retrieval head takes the normalized 1,024-dimensional TM-head vector as input (Supplementary Fig. 4c). It applies layer normalization^53^, Linear(1,024, 1,024), GELU^57^ and Linear(1,024, 1,024), followed by a residual connection and L2 normalization (2.10 million parameters). The final layer is zero-initialized, and the encoder and TM head stay frozen. The head is trained with a Euclidean triplet margin loss^59^ (margin 1.0) on same-EC positives and different-EC negatives, using Adam^60^ (learning rate 10⁻⁴; no weight decay), 2,048 triplets per step and 4,000 steps.

Training data follow the DEER curation^49^. The CARE enzyme dataset^61^, reconstructed from Swiss-Prot^54^ (release 2024.05.13), is split by EC class into 62,350 training and 3,343 validation proteins. The checkpoint is selected on the validation set.

### Residue-level alignment

The alignment head is a residual perceptron that acts directly on frozen residue features, bypassing the TM head (Supplementary Fig. 4c). It applies Linear(1,024, 2,048), GELU^57^ and Linear(2,048, 1,024), followed by a residual connection and per-residue L2 normalization (4.20 million parameters). The final layer is zero-initialized.

Following You and colleagues^37^, the head is trained with residue-level InfoNCE (temperature 0.07) on correspondences from structural alignments, using the released 10,000 training pairs and 300 validation pairs for checkpoint selection. Training runs for 60 epochs on cached residue features with 128 pairs per batch, using Adam^60^ (peak learning rate 2 × 10⁻³; cosine decay; 5% warm-up; weight decay 0.1).

Query–target inner products of the head outputs form a cosine score matrix that is decoded by global Needleman–Wunsch alignment with gap penalty 0, using the reference implementation^37^ (Supplementary Fig. 4c, d). Alignment performance is measured as the F1 over all aligned residue pairs, relative to curated structural alignments. We report results separately for two datasets: MALIDUP^50^, which contains 241 pairs of homologous duplicated domains (median sequence identity 13.8%), and MALISAM^40^, which contains 130 pairs of structurally analogous but non-homologous proteins (median sequence identity 8.0%).

### Pfam-clan candidate prefilter

The Pfam-clan prefilter is adapted from PLMSearch^25^. PfamScan assigns Pfam families to each query and target, and each family is mapped to its clan, or kept as its own identifier when it has no clan. A target is a candidate when it shares an identifier with the query. If no target does, the query searches the full database. Targets outside the candidate set are excluded from ranking, and the scores of the others are unchanged. The filter adds no learned parameters.

### Sequence-only remote homology search

Query and database sequences pass through the frozen encoder, mean pooling and the TM head, and EC-specific retrieval additionally applies the EC head (Supplementary Fig. 4d). Query and database sequences are encoded at full length without truncation. Their 1,024-dimensional vectors are computed once and cached, and each query is scored against all database vectors by cosine similarity, which for normalized EC vectors gives the same ranking as Euclidean distance. For TM-head vectors *z*q and *z*t, an optional TM-score estimate uses the scale *a* and offset *b* learned with the TM head (Equation (11)):

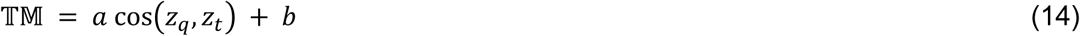

Ranking uses the cosine scores directly, and Equation (14) applies only to TM-head vectors. The optional Pfam-clan prefilter is applied after scoring and removes non-candidate targets from the ranking.

For selected hits, the alignment head embeds query and target residues, and their *L*q × *L*t cosine matrix is aligned by global Needleman–Wunsch with zero gap penalty^37^ (Supplementary Fig. 4d). Other dynamic-programming aligners, such as local Smith– Waterman alignment, can be applied to the same matrix but were not used here.

Inference needs no structures, EMA branches or training projectors.

### Remote-homology retrieval on SCOPe

We used the retrieval benchmark released with DHR^27^, built on SCOPe 2.08^38^. One representative domain of each family or superfamily is searched against all 29,421 SCOPe domains. A hit is correct when it shares the query’s family or, at the second level, its superfamily. Query–target pairs are stratified into ten bins of sequence identity, and results are reported per bin. Following the benchmark, AUC-1FP is the fraction of the query’s true positives retrieved before the first false positive, averaged over queries within each identity bin. Top-*K* recall is the fraction of true relatives found within the first *K* hits. S-PLM-TM uses the Pfam-clan prefilter, as does PLMSearch, and methods without a prefilter search the full database. Error bars are standard error of the mean (s.e.m.) over queries.

### Fold classification on CATH

We used the fold-classification task of a recent benchmark of remote-homology search methods^47^. Each of the 15,043 CATH-S20^39^ domains is searched against all others, and a hit is correct when it shares the query’s class, architecture, topology or homologous-superfamily label at the level being scored. The main metric is AUC-1FP, defined as for SCOPe and computed from each method’s output with the benchmark script. F1-max, the maximum F1 over all score thresholds (Supplementary Fig. 2), follows the benchmark convention, in which a domain with no true positives at a given classification level is assigned a score of zero.

### Functional consistency by Gene Ontology

We used the functional-consistency task of the same benchmark^47^. The top 10 targets retrieved for each query are scored against the query by Gene Ontology^48^ molecular-function semantic similarity with a best-match average. Results are reported for all annotated pairs and for the experimentally supported subset (evidence codes EXP, IDA, IMP, IPI, TAS and IC), with one annotation file and one scorer for all methods. No Pfam-clan prefilter is applied to S-PLM-TM in this benchmark. Baselines are run with their default pipelines. Descriptive means and s.e.m. are computed over annotated query– target pairs, whereas significance tests are paired by query (see Statistical testing). The experimentally supported subset contains 150 queries with qualifying annotations on both sides.

### Microbial–host isozyme retrieval

We used the isozyme benchmark released with DEER^49^, in which 90 human enzyme templates, one per EC family, are searched against 4,213 bacterial enzymes containing 449 true isozymes. A retrieved enzyme is correct when it carries the template’s EC number. S-PLM-TM ranks targets by cosine similarity between TM-head vectors, and the EC-specific model by Euclidean distance between EC-head vectors (Supplementary Fig. 4d). Metrics follow the DEER definitions. For a template with *n* true counterparts, recall is the fraction recovered within its top *n* hits, averaged over templates. At cut-offs *k* = 10, 30 and 100, hit ratio is the fraction of the 449 isozymes retrieved by at least one template within its top *k* hits, and relative precision is the number of correct hits in the top *k* divided by min(*k*, *n*), averaged over templates. Error bars are s.e.m. over the 90 templates.

### Search at scale

To search at scale, the target library was encoded once and cached. Library sequences were sorted by length so that sequences of similar length fell in the same batch. A batch held at most eight sequences and at most 8,192 padded tokens, counted as the number of sequences times the length of the longest sequence in the batch. Sequences were not truncated, and padding positions were masked during pooling. The resulting TM-head vectors were stored as a single matrix. Targets were then ranked by the cosine similarity between the query vector and this matrix, which costs one matrix multiplication per query. Queries are encoded by the same procedure and can be batched in the same way, but this option was not used here: each query was encoded individually in all the benchmarks reported here.

We assembled a large-scale benchmark in which SCOPe40^38^ domains are searched against a Swiss-Prot library. Its query and target sets were constructed here from the sequence collections released with PLMSearch^25^ and are not those of that benchmark. The target library consists of 430,140 Swiss-Prot^54^ proteins with an AlphaFold DB^14^ structure, from which proteins with a mean pLDDT below 70 or with at least 40% sequence identity to SCOPe40-train have been removed. Candidate queries were taken from SCOPe40-test, which shares no fold with SCOPe40-train, capped at 30 per superfamily and shuffled with a fixed seed, and each candidate was aligned against every protein in the library with TM-align^24^. A candidate was retained when at least ten library proteins reached a TM-score above 0.5 at below 30% sequence identity to it, which left 957 query domains from 242 superfamilies.

The ground truth is this same all-against-all TM-align^24^ matrix, which we computed ourselves rather than reusing the similarity matrix distributed with PLMSearch^25^. TM-scores were normalized by the average length of the two structures, and a pair counts as positive when its TM-score exceeds 0.5. Pairs whose lengths differ at least three-fold cannot exceed a TM-score of 0.5 under this normalization and were therefore treated as negatives without being aligned. The resulting matrix contains 708,472 positive pairs. Each query is excluded from its own ranking. Results are reported over all positive pairs and over the subset below 30% sequence identity to the query, where sequence identity is the fraction of identical residues over all columns of a Needleman–Wunsch global alignment computed with the BLOSUM62 matrix and gap opening and extension penalties of 10 and 0.5. Performance is reported as mean average precision (MAP), the mean over queries of the average precision of each query’s ranking, and as precision at ranks 1 and 10 (P@1 and P@10). For evaluation by sequence identity, positive targets were grouped into 10-percentage-point bins according to their identity to each query. Within each bin, positives outside the bin were removed from the ranking, while all negatives (TM-score ≤ 0.5) were retained. MAP was averaged only over queries with at least one positive in that bin; a query could therefore contribute to multiple bins.

### Baseline methods

Baselines fall into three groups: alignment-based structure comparison (TM-align^24^, Dali^29^, GTalign^30^ and Foldseek^31^ with its TM-score mode), representation-based structure comparison (GraSR^32^ and FoldExplorer^33^), and sequence-based search. The last group comprises representation-based methods (TM-Vec^23^, PLMSearch^25^ and its SS-predictor, DHR^27^, ERAST^28^) and alignment-based tools (BLASTp and PSI-BLAST^1^, DIAMOND^5^, MMseqs2^4^, phmmer and jackhmmer^2^, and Foldseek in ProstT5 mode^3^). Baseline results in the retrieval benchmarks were generated in house from released weights or binaries, using the same labels, annotation files and scoring code as S-PLM-TM; residue-alignment comparator values in Supplementary Table 2 were taken from CLAlign under its reported protocol. Foldseek in ProstT5 mode was run with a cap of 2,000 hits per query, which limits its recall on the large-scale benchmark.

### Statistical testing and multiple-testing correction

Differences from S-PLM-TM were assessed using two-sided Wilcoxon signed-rank tests, paired per query for SCOPe, Gene Ontology and the large-scale benchmark, per domain for CATH, and per EC-family template for bacterial EC retrieval. *P* values were adjusted for multiple comparisons using the Benjamini–Hochberg procedure and are reported as adjusted *P* values. Each correction was applied within the set of comparisons reported together, that is, the cells of the table or figure panel in which a comparison appears, and each such family was fixed before the tests were run. Where a group of comparisons is summarized jointly, the largest adjusted *P* value in the group is reported as an upper bound.

### Timing protocol

Timings were measured on the same node type: CPU-only methods used 16 CPU cores, and GPU-enabled methods used one A100 80 GB GPU. Search time includes query encoding or preparation, database scoring and result writing, but excludes model loading and one-time target-index construction; index time is reported separately. Vector-based methods used exhaustive scoring against the target database. TM-align and Dali were timed on 100 queries against a fixed random 1% sample of targets, and their times were extrapolated to the full benchmark by query and target counts. These times are marked as extrapolated in Table 1. Their accuracy values come from separate runs on the full benchmark.

### Ablation studies

Two control encoders were pretrained to assess the cross-modal objective. The first omitted flow matching at both representation levels and in both cross-modal directions. The second replaced flow matching with a mean-squared-error (MSE) regression loss between the sequence and structure projections at the residue and protein levels (weight 1.0). All other objectives, data and training settings were unchanged. Each control received its own TM-score calibration head and the same evaluation as S-PLM-TM (Supplementary Table 4).

## Data availability

The S-PLM-TM model weights, including the sequence encoder, TM-score calibration head, EC-number retrieval head and residue-level alignment head, are available at https://huggingface.co/deepduoduo/S-PLM-TM. Benchmark data for the SCOPe and large-scale retrieval evaluations, including the query and target sets used, annotations, reference scores and per-query evaluation results, are available at https://huggingface.co/datasets/deepduoduo/S-PLM-TM-benchmarks. The SCOPe query and target sets were derived from the public DHR release and its authors; the exact sets used here are redistributed in that dataset. Target protein structures for the large-scale benchmark are not redistributed but can be retrieved from the AlphaFold Protein Structure Database using the provided accessions.

## Code availability

Code for generating embeddings, retrieval scores and residue-level alignment scores is available at https://github.com/duolinwang/S-PLM-TM.

## Acknowledgements

This work was supported by the National Institutes of Health (NIH) under grant numbers R01-LM014510, R01-LM015024, and R35-GM126985. We thank Dr. Jin Chen at the University of Alabama at Birmingham for his valuable comments and advice on this work.

## Contributions

D.X. and D.W. conceived the project and designed the experiments. D.W. and Y.Z. performed the experiments and analyses, with input from S.Z. and Q.S. Q.S. advised on the project. D.W. wrote the manuscript, with input from Y.Z., S.Z., Q.S. and D.X.

