## Supplemental materials for "S-PLM-TM enables fast and accurate sequence-based search for remote homologs"

### Supplementary Information

#### Supplementary Figures

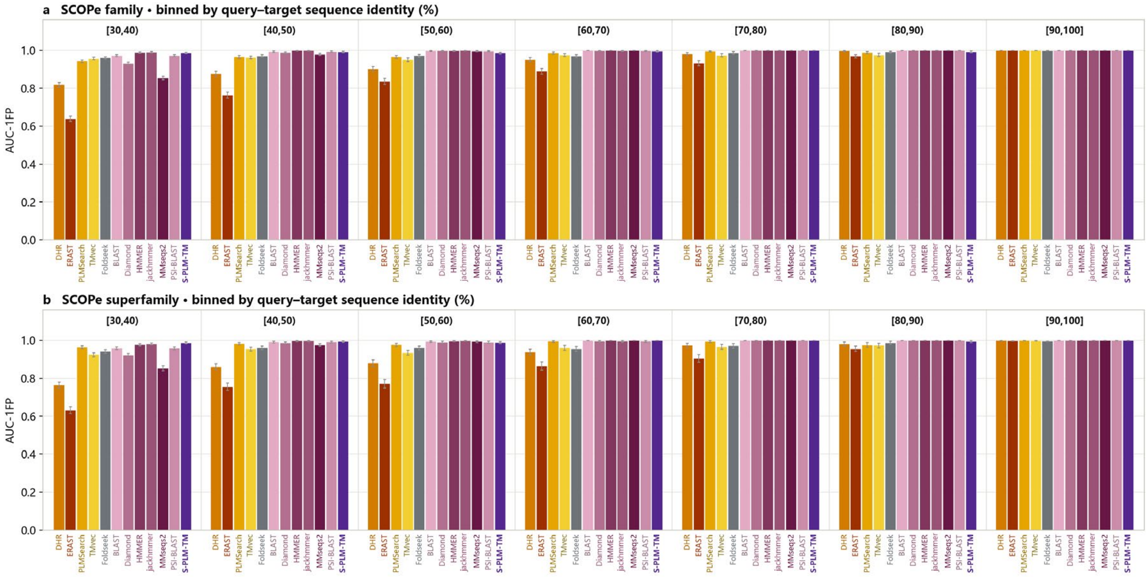


**Supplementary Fig. 1 | SCOPe homology detection at ≥30% sequence identity.** AUC-1FP for **a**, family-level and **b**, superfamily-level retrieval, binned by query–target sequence identity. The three bins below 30% identity are shown in Fig. 2ab. Bars are means ± s.e.m. over queries.

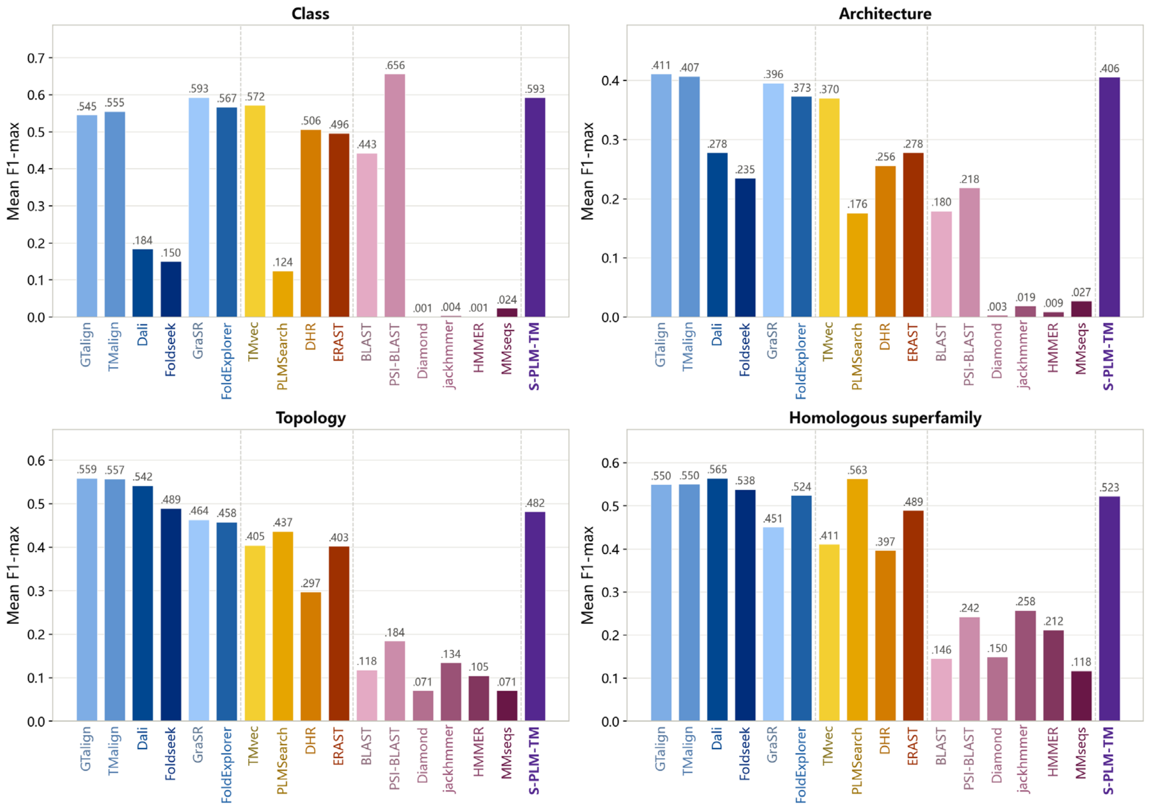


**Supplementary Fig. 2. CATH fold classification scored as mean F1-max.** Mean F1-max at each of the four CATH levels, computed from each method's ranked output. Following the published benchmark, a domain with no true positive at a level is scored 0, whereas a query for which a method returns no hits is excluded from that method's mean. Our reproduced baselines match the published values (mean absolute difference 0.007 over 56 values); the exception is PLMSearch, which deviates at three levels: our reproduction scores 0.563 versus 0.51 published at homologous superfamily and 0.437 versus 0.41 at topology, but 0.124 versus 0.16 at class. Its Pfam pre-filter returns a short candidate list, which raises F1-max at homologous superfamily, where a query has only a few true positives, and lowers it at class level, where a query can have thousands of true positives and a short list caps the attainable recall.

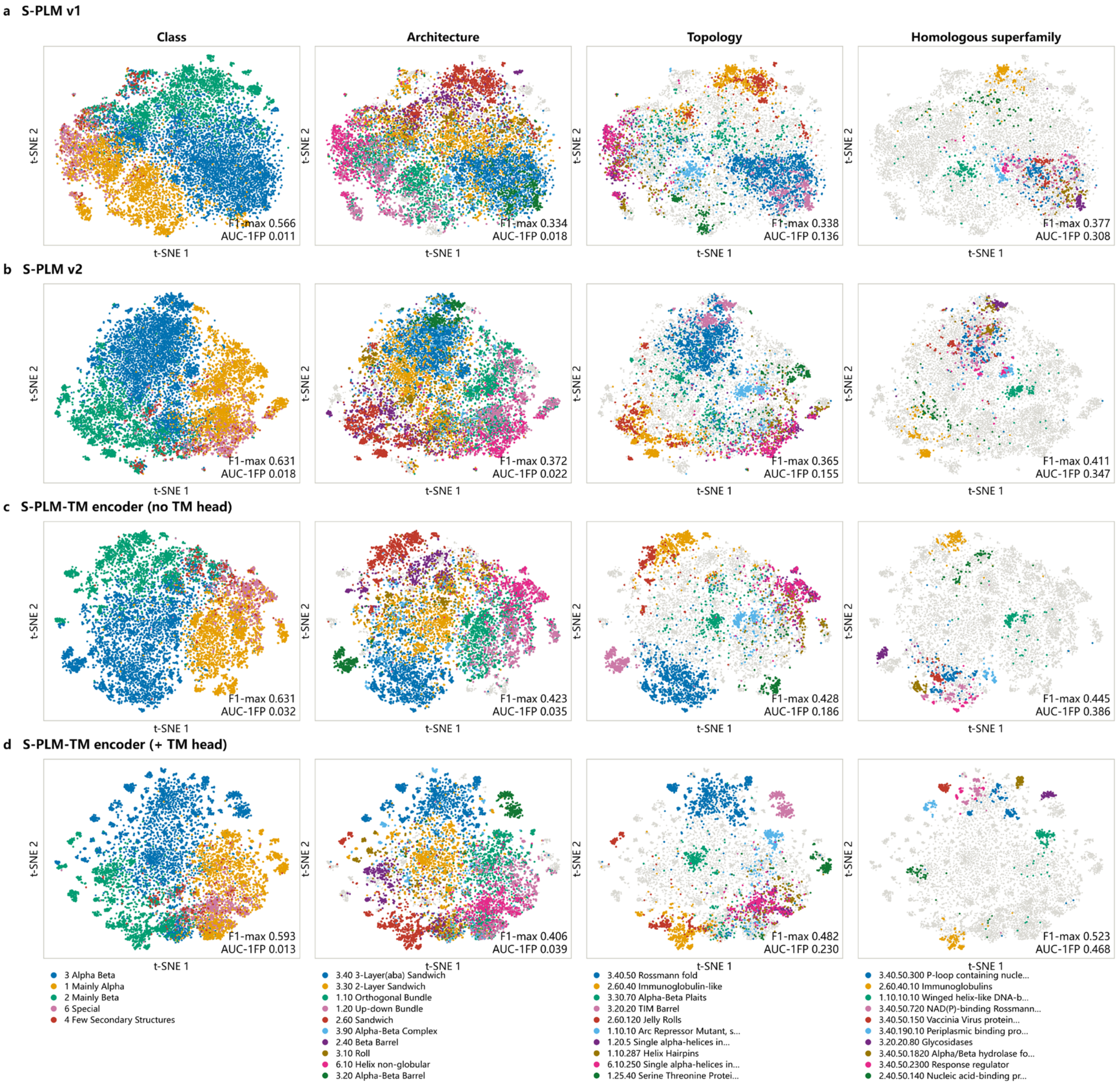


**Supplementary Fig. 3. Evolution of the CATH-S20 embedding space across S-PLM versions.** Two-dimensional t-SNE projections of the sequence embeddings of all 15,043 CATH-S20 domains, for **a**, S-PLM v1; **b**, S-PLM v2; **c,** the S-PLM-TM sequence encoder before the TM-score calibration head; and **d**, the same encoder after the head. Within each row the four panels show the same projection colored by the largest categories at the class (all five classes), architecture, topology and homologous-superfamily levels (the ten largest categories at these levels); domains outside those categories are grey. Mean F1-max and AUC-1FP for the corresponding model and level are given in each panel.

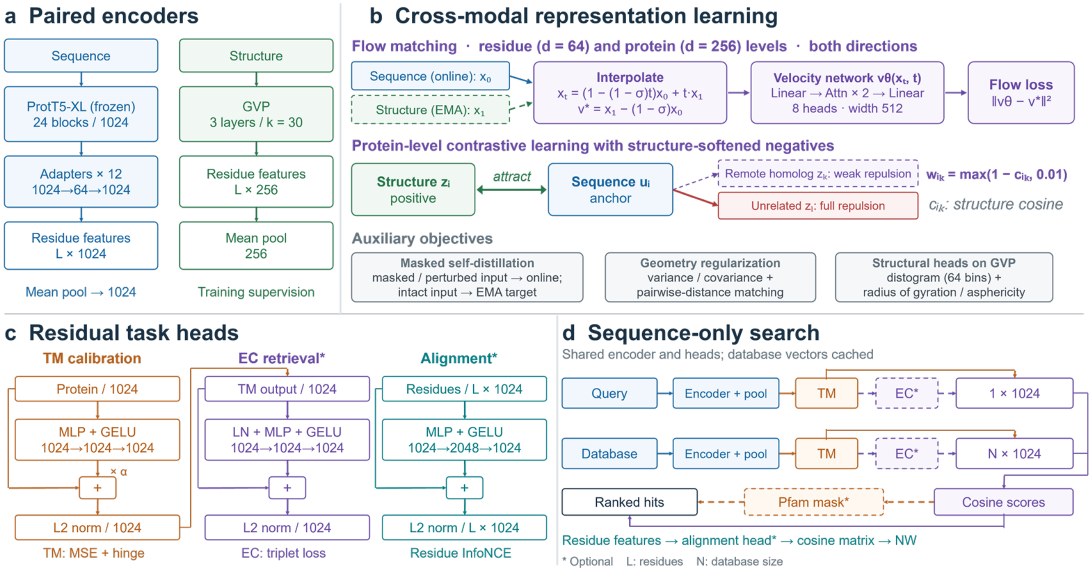


**Supplementary Fig. 4. Architecture and workflow of S-PLM-TM.** **a**, Paired sequence and structure encoders. **b**, Cross-modal representation learning. **c**, Residual heads for TM-score calibration, EC retrieval and residue alignment. **d**, Sequence-only remote homology search.

### Supplementary Tables

##### Supplementary Table 1. Search performance by sequence identity to the query.

| Method | Input | MAP | | |
|---|---|---|---|---|
| | | 0–10% (307 queries) | 10–20% (950 queries) | 20–30% (876 queries) |
| TM-align | St | 1.000 | 1.000 | 1.000 |
| GTalign | St | 0.806 | 0.968 | 0.947 |
| **S-PLM-TM** | **S** | 0.125 | **0.569** | **0.665** |
| TM-Vec | S | **0.136** | 0.532 | 0.600 |
| FoldExplorer | St | 0.109 | 0.522 | 0.609 |
| PLMSearch | S | 0.108 | 0.518 | 0.587 |
| Dali | St | 0.077 | 0.487 | 0.527 |
| GraSR | St | 0.173 | 0.472 | 0.556 |
| SS-predictor | S | 0.083 | 0.416 | 0.508 |
| Foldseek (ProstT5) | S | 0.062 | 0.381 | 0.507 |
| DHR | S | 0.053 | 0.373 | 0.506 |
| Foldseek-TM | St | 0.077 | 0.356 | 0.472 |
| Foldseek | St | 0.037 | 0.321 | 0.357 |
| jackhmmer | S | 0.040 | 0.297 | 0.462 |
| PSI-BLAST | S | 0.039 | 0.270 | 0.451 |
| phmmer | S | 0.017 | 0.162 | 0.418 |
| BLASTp | S | 0.012 | 0.134 | 0.381 |
| MMseqs2 | S | 0.007 | 0.128 | 0.369 |
| DIAMOND | S | 0.008 | 0.130 | 0.366 |
| ERAST | S | 0.023 | 0.118 | 0.261 |

Input: S, sequence; St, structure. MAP, mean average precision. Sequence-identity bins are defined for query–positive-target pairs. Counts in the header indicate queries with at least one positive in each bin and overlap across bins. TM-align defines the benchmark ground truth and therefore scores 1.000 by construction; it is included for reference only. The query pool comprises the 957 SCOPe domains used in the large-scale benchmark. S-PLM-TM was run without the Pfam-clan prefilter.

##### Supplementary Table 2. Residue-level structural alignment.

| Method | Trainable parameters | MALIDUP F1 | MALISAM F1 |
|---|---|---|---|
| S-PLM-TM encoder + alignment head | 4.20 M | 0.643 | 0.282 |
| CLAlign (ProstT5, LoRA r = 8) | 10.62 M | **0.677** | **0.299** |
| CLAlign (ProtT5, LoRA r = 8) | 10.62 M | 0.646 | 0.267 |
| PLMAlign | — | 0.561 | 0.184 |
| TEA-NW | — | 0.556 | 0.176 |
| DeepBLAST | — | 0.518 | 0.152 |
| DEDAL | — | 0.417 | 0.070 |

Full-alignment F1 was evaluated against hand-curated structural alignments using CLAlign's Needleman–Wunsch implementation with zero gap penalty[37]. MALIDUP[50] contains 241 homologous internal-duplication pairs, whereas MALISAM[40] contains 130 structurally analogous, non-homologous pairs. All comparator results were taken from CLAlign under the same protocol; its strongest variant uses a LoRA-adapted 3-billion-parameter ProstT5 backbone.

##### Supplementary Table 3. Encoder ablation across protein language models.

| Benchmark and metric | S-PLM-TM | ProtT5-XL | ProstT5 | ESM2-650M |
|---|---|---|---|---|
| **SCOPe, AUC-1FP, family level** | | | | |
| 0–10% identity | **0.482** | **0.482** | 0.459 | 0.466 |
| 10–20% | **0.788** | 0.769 | 0.781 | 0.745 |
| 20–30% | 0.942 | 0.934 | **0.946** | 0.928 |
| **SCOPe, AUC-1FP, superfamily level** | | | | |
| 0–10% identity | **0.560** | 0.537 | 0.542 | 0.524 |
| 10–20% | **0.763** | 0.742 | 0.757 | 0.721 |

| Benchmark and metric | S-PLM-TM | ProtT5-XL | ProstT5 | ESM2-650M |
|---|---|---|---|---|
| 20–30% | 0.926 | 0.915 | **0.928** | 0.903 |
| **CATH fold classification, F1max** | | | | |
| Class | **0.593** | 0.552 | 0.569 | 0.558 |
| Architecture | **0.406** | 0.334 | 0.374 | 0.336 |
| Topology | **0.482** | 0.397 | 0.455 | 0.379 |
| Homologous superfamily | **0.523** | 0.478 | 0.513 | 0.443 |
| **CATH fold classification, AUC-1FP** | | | | |
| Class | **0.013** | 0.006 | 0.010 | 0.008 |
| Architecture | **0.039** | 0.023 | 0.033 | 0.022 |
| Topology | **0.230** | 0.178 | 0.214 | 0.165 |
| Homologous superfamily | **0.468** | 0.401 | 0.461 | 0.367 |
| **GO molecular function, semantic similarity (experimentally supported subset)** | | | | |
| Mean | **0.492** | 0.482 | 0.470 | 0.456 |
| Median | **0.507** | 0.484 | 0.482 | 0.429 |
| **Bacterial EC retrieval** | | | | |
| Mean recall | **0.266** | 0.223 | 0.216 | 0.239 |
| Hit ratio at 10 | **0.303** | 0.276 | 0.261 | 0.267 |
| Relative precision at 10 | **0.318** | 0.304 | 0.288 | 0.291 |
| Hit ratio at 30 | **0.463** | 0.432 | 0.430 | 0.423 |
| Relative precision at 30 | **0.378** | 0.352 | 0.363 | 0.341 |
| Hit ratio at 100 | 0.572 | 0.564 | 0.566 | **0.584** |
| Relative precision at 100 | 0.491 | 0.490 | 0.491 | **0.498** |

All encoders are frozen and paired with identically trained TM-score calibration heads. Evaluation settings and metrics follow the main text. Pfam filtering is applied to all evaluations except CATH and GO. Bold indicates the best result in each row.

##### Supplementary Table 4. Component ablation of S-PLM-TM.

| Benchmark and metric | S-PLM-TM | No flow matching | MSE regression | No TM-head | No Pfam prefilter | Raw encoder |
|---|---|---|---|---|---|---|
| **SCOPe, AUC-1FP, family level** | | | | | | |
| 0–10% identity | **0.4816** | 0.4808 | 0.4760 | 0.4308 | 0.2246 | 0.0753 |
| 10–20% | **0.7879** | 0.7853 | 0.7819 | 0.7407 | 0.7136 | 0.5771 |
| 20–30% | 0.9424 | **0.9439** | 0.9427 | 0.9291 | 0.9229 | 0.8845 |
| **SCOPe, AUC-1FP, superfamily level** | | | | | | |
| 0–10% identity | **0.5598** | 0.5556 | 0.5524 | 0.4953 | 0.1747 | 0.0599 |
| 10–20% | 0.7629 | 0.7632 | **0.7644** | 0.7450 | 0.6183 | 0.4871 |
| 20–30% | **0.9264** | 0.9226 | 0.9249 | 0.9192 | 0.8742 | 0.8301 |
| **CATH fold classification, F1max** | | | | | | |
| Class | 0.5927* | 0.5917 | 0.5961 | **0.6309^** | 0.5927* | **0.6309^** |
| Architecture | 0.4057* | 0.3975 | 0.4074 | **0.4226^** | 0.4057* | **0.4226^** |
| Topology | **0.4815*** | 0.4725 | 0.4749 | 0.4278^ | **0.4815*** | 0.4278^ |
| Homologous superfamily | **0.5228*** | 0.5200 | 0.5159 | 0.4451^ | **0.5228*** | 0.4451^ |
| **CATH fold classification, AUC-1FP** | | | | | | |
| Class | 0.0135* | 0.0128 | 0.0143 | **0.0321^** | 0.0135* | **0.0321^** |
| Architecture | **0.0394*** | 0.0370 | 0.0391 | 0.0354^ | **0.0394*** | 0.0354^ |
| Topology | **0.2300*** | 0.2257 | 0.2262 | 0.1857^ | **0.2300*** | 0.1857^ |
| Homologous superfamily | **0.4678*** | 0.4620 | 0.4595 | 0.3855^ | **0.4678*** | 0.3855^ |
| **GO molecular function, semantic similarity (experimentally supported subset)** | | | | | | |
| Mean | **0.4915*** | 0.4882 | 0.4777 | 0.4402^ | **0.4915*** | 0.4402^ |
| Median | **0.5067*** | 0.5067 | 0.4867 | 0.3987^ | **0.5067*** | 0.3987^ |
| **Bacterial EC retrieval** | | | | | | |
| Mean recall | **0.2659** | 0.2540 | 0.2306 | 0.1820 | 0.2450 | 0.1295 |
| Hit ratio at 10 | **0.3029** | 0.3007 | 0.2829 | 0.2339 | 0.2784 | 0.1737 |
| Relative precision at 10 | **0.3177** | 0.3119 | 0.3018 | 0.2612 | 0.2957 | 0.1762 |
| Hit ratio at 30 | **0.4633** | 0.4610 | 0.4521 | 0.4143 | 0.4365 | 0.2873 |
| Relative precision at 30 | **0.3781** | 0.3686 | 0.3729 | 0.3527 | 0.3624 | 0.2242 |
| Hit ratio at 100 | **0.5724** | 0.5702 | 0.5657 | 0.5457 | 0.5434 | 0.3742 |
| Relative precision at 100 | **0.4906** | 0.4896 | 0.4868 | 0.4763 | 0.4245 | 0.3091 |

The first three columns compare pretraining objectives under otherwise identical training settings, including TM-score refinement and separately trained calibration heads. No flow matching removes all the flow-matching losses; MSE regression replaces them with MSE losses between sequence and structure projections at the residue and protein levels. The last three columns use the same S-PLM-TM encoder without retraining: No TM-head removes only the TM-score calibration head; No Pfam prefilter removes the Pfam-clan candidate filter; Raw encoder removes both. Evaluation settings and metrics follow the main text. CATH and GO use no Pfam prefilter, so columns 1 and 5 (*) are identical, as are columns 4 and 6 (^). Bold indicates the best result in each row.

#### Supplementary Table 5. Effect of the Pfam-clan prefilter in the large-scale search benchmark.

| Metric or identity bin | Number of queries | S-PLM-TM | S-PLM-TM + Pfam-clan prefilter |
|---|---|---|---|
| a, Overall performance | | | |
| MAP | 957 | **0.638** | 0.601 |
| MAP (<30%) | 957 | **0.620** | 0.584 |
| P@1 | 957 | 0.844 | **0.861** |
| P@10 | 957 | **0.805** | 0.801 |
| b, MAP by sequence identity of the positive targets to the query | | | |
| 0–10% | 307 | **0.125** | 0.110 |
| 10–20% | 950 | **0.569** | 0.543 |
| 20–30% | 876 | **0.665** | 0.647 |
| 30–40% | 465 | **0.912** | **0.912** |
| 40–50% | 323 | 0.905 | **0.937** |
| 50–60% | 239 | 0.931 | **0.957** |
| 60–70% | 180 | 0.953 | **0.973** |
| 70–80% | 159 | 0.990 | **0.994** |
| 80–90% | 153 | 0.993 | **1.000** |
| 90–100% | 224 | **1.000** | **1.000** |

Table 1 and Supplementary Table 1 report S-PLM-TM without the Pfam-clan prefilter. Here the prefilter is added back. Overall metrics use the same 957 queries as Table 1; each sequence-identity bin includes only queries with at least one positive in that bin, so query counts overlap across bins. Bold indicates the higher of the two values in each row, or both when they are equal.
